# PRECISE: A domain adaptation approach to transfer predictors of drug response from pre-clinical models to tumors

**DOI:** 10.1101/536797

**Authors:** Soufiane Mourragui, Marco Loog, Marcel JT Reinders, Lodewyk FA Wessels

## Abstract

**Motivation:** Cell lines and patient-derived xenografts (PDX) have been used extensively to understand the molecular underpinnings of cancer. While core biological processes are typically conserved, these models also show important differences compared to human tumors, hampering the translation of findings from pre-clinical models to the human setting. In particular, employing drug response predictors generated on data derived from pre-clinical models to predict patient response, remains a challenging task. As very large drug response datasets have been collected for pre-clinical models, and patient drug response data is often lacking, there is an urgent need for methods that efficiently transfer drug response predictors from pre-clinical models to the human setting.

**Results:** We show that cell lines and PDXs share common characteristics and processes with human tumors. We quantify this similarity and show that a regression model cannot simply be trained on cell lines or PDXs and then applied on tumors. We developed PRECISE, a novel methodology based on domain adaptation that captures the common information shared amongst pre-clinical models and human tumors in a consensus representation. Employing this representation, we train predictors of drug response on pre-clinical data and apply these predictors to stratify human tumors. We show that the resulting domain-invariant predictors show a small reduction in predictive performance in the pre-clinical domain but, importantly, reliably recover known associations between independent biomarkers and their companion drugs on human tumors.

**Availability:** PRECISE and the scripts for running our experiments are available on our GitHub page (https://github.com/NKI-CCB/PRECISE).

**Supplementary information:** Supplementary data are available. online.

## 1 Introduction

Cancer is a heterogeneous disease that arises due to the accumulation of somatic genomic alterations. These alterations show high levels of variability between tumors resulting in heterogeneous responses to treatments. Precision medicine attempts to improve response rates by taking this heterogeneity into account and tailoring treatment to the specific molecular make-up of a given tumor. This requires the identification of biomarkers to identify the set of patients that will benefit from a given treatment while sparing those that will not benefit the unnecessary side-effects. However, as there are limited patient response data for a wide range of drugs, pre-clinical modes such as cell lines and patient-derived xenografts (PDXs) have been employed to generate large-scale data sets that enable the development of personalized treatment strategies based on the data-driven identification of biomarkers of response. More specifically, hundreds of pre-clinical models have not only been extensively molecularly characterized, but, more importantly, their response to hundreds of drugs have also been recorded. This has resulted in large public resources containing data derived from cell lines (GDSC1000, Iorio *et al.* [2016]) and PDX models (NIBR PDXE, Gao *et al.* [2015]).

These pre-clinical resources can be employed to build predictors of drug response which are then transferred to the human setting, allowing stratification of patients for drugs the patients have not yet been exposed to. Geeleher *et al.* applied this approach by simply correcting for a batch effect between the cell line and tumor data sets and then directly transferring the cell line predictor to the human setting (Geeleher *et al.* [2014, 2017]). This already yielded some promising results: it recovered well-established biomarkers such as the association between Lapatinib sensitivity and ERBB2 amplifications. However, when directly transferring a predictor from the source domain (cell lines) to the target domain (human tumors) one assumes that the source and target data originate from the same distribution. While the differences between pre-clinical models and human tumors have been studied extensively (Gillet *et al.* [2013], Ben-David *et al.* [2018], Ben-David *et al.* [2017]), the most obvious differences include the absence of an immune system in both cell lines and PDXs and the absence of a tumor micro-environment and vasculature in cell lines. One can therefore not assume similarity between the source and target distributions.

Transfer Learning aims at addressing this issue (see (Pan and Yang [2010]) for a general review). Transfer learning methods can be assigned to different categories depending on the availability of source and target labels and on the specific relation between these source and target datasets. Since we have a vey small number of labeled tumor samples, but a wealth of labeled pre-clinical models, our approach falls into the category referred to as *transductive*^1^. Since the features (i.e. the genes) are the same in the source and target domains, our problem requires a *domain adaptation* strategy, sometimes also referred to as *homogeneous domain adaptation*.

As previously mentioned, the marginal distributions of pre-clinical models and tumors are expected to be different. However, we assume that drug response is, for a large part, determined by biological phenomena that are conserved between pre-clinical models and human tumors. Therefore there should exist a set of features (genes) for which the conditional distribution of drug response given these features is comparable across cell lines, PDXs and human tumors. Different methodologies have been proposed to find such a common space and these can be divided in two main categories (Csurka [2017]). In the first approach, a common subspace can be found directly from both the pre-clinical models and the tumors by aligning the marginal distributions. This can be done by, for instance, using the Maximum Mean Discrepancy either exactly or employing semi-definite-programming (Song *et al.* [2008], Pan *et al.* [2008]), or by using approximations based on multiple-kernel learning (Duan *et al.* [2012]) or empirical kernel maps (Pan *et al.* [2011]). A second way is to perform the domain adaptation correction by first reducing the dimensionality, and then aligning the low-rank representations (Fernando *et al.* [2013], Gopalan *et al.* [2011], Gong *et al.* [2012]). Approaches in the second category are more robust against small sample sizes as they do not try to match distributions exactly, but instead find the most important (lower dimensional) directions in each dataset and map these onto each other. We therefore chose to employ the latter approach to transfer response predictors from pre-clinical models to human tumors.

We present PRECISE (**P**atient **R**esponse **E**stimation **C**orrected by **I**nterpolation of **S**ubspace **E**mbeddings), a methodology based on domain adaptation that trains a regression model on processes that human tumors share with pre-clinical models. Fig 1 shows the general workflow of PRECISE. We first independently extract factors from the cell lines, PDXs and human tumors by means of linear dimensionality reduction. We then use a linear transformation that geometrically matches the factors from one of the pre-clinical models to the human tumor factors (Fernando *et al.* [2013]). Subsequently we extract the common factors (*principal vectors (PVs)*) defined as the directions that are the least influenced by the linear transformation (Fig 1A). After selection of the most similar principal vectors, we compute new feature spaces (based on this selection) by interpolating between the source domain (cell line or PDX principal vectors) and the target domain (human tumor principal vectors). The feature spaces resulting from this interpolation allow a balance to be struck between the chosen model system and the tumors (Gopalan *et al.* [2011], Gong *et al.* [2012]).^2^ From the set of interpolated spaces, the *consensus representation* is obtained by optimizing the match between the marginal distributions of the chosen pre-clinical model and the human tumor data projected on these interpolated features. These consensus features are finally used to train a regression model using data from the pre-clinical model of choice. We use this regression model to predict tumor drug response (Fig 1B). As these features are shared between the pre-clinical models and the human tumors, the regression model is expected to generalize to human tumors. We finally use known biomarker-drug associations (from independent data sources, e.g. mutation status, copy number) as positive controls to validate the predictions of the model in human tumors (Fig 1C).

**Fig. 1.**
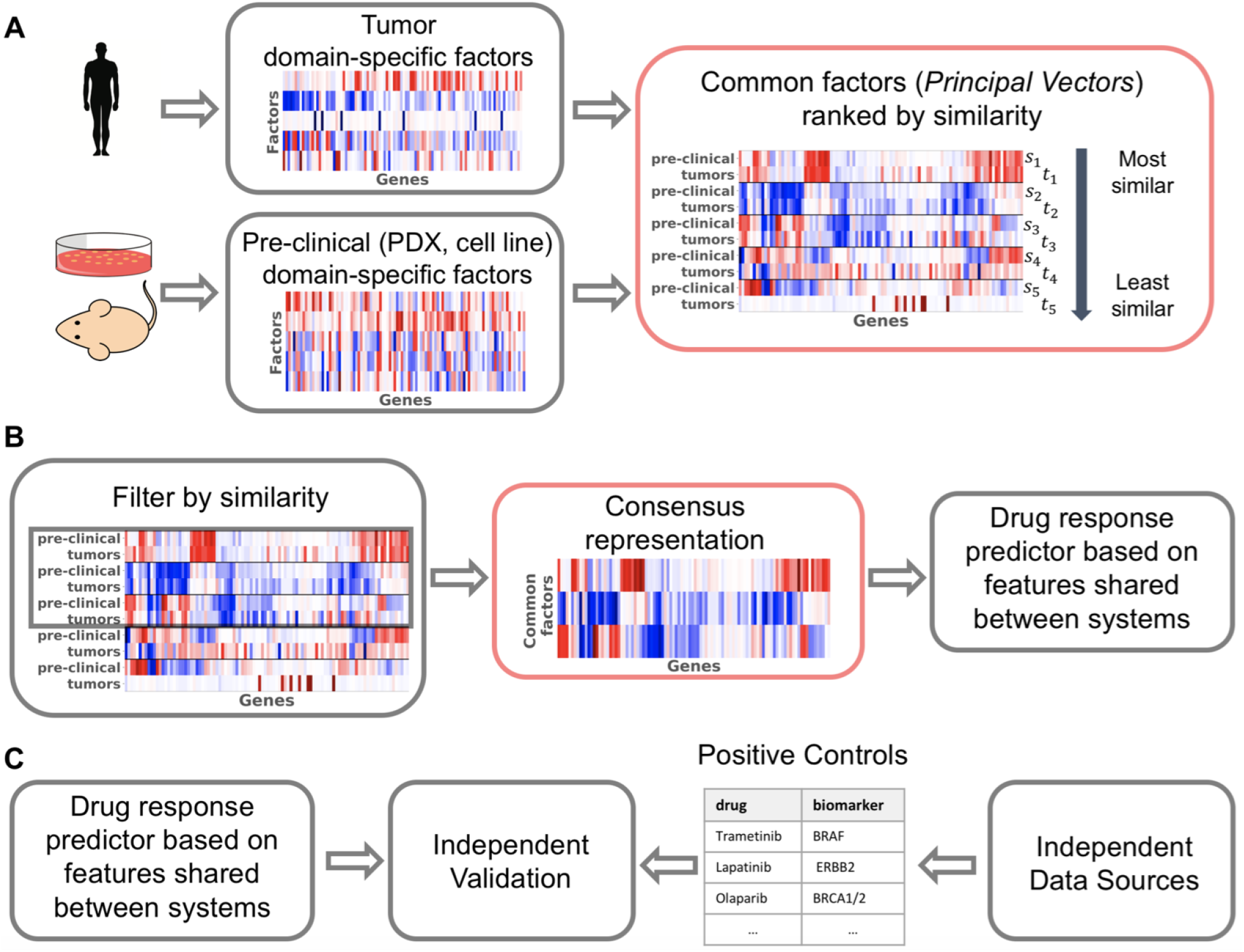
Overview of PRECISE and its validation. **(A)** Human tumor and pre-clinical data are first processed independently to find the most important domain-specific factors (using, for instance PCA). These factors are then compared, aligned and ordered by similarity, yielding principal vectors (PVs). The first PVs are pairs of vectors that are geometrically very similar and capture strong commonality between human tumors and pre-clinical models, the PVs at the bottom represent dissimilarities between human tumors and pre-clinical models. **(B)** A cut-off in similarity enables the retention of processes that are common. After interpolation between these most similar pre-clinical and tumor PVs, a consensus representation is computed by balancing the influence of human tumor and pre-clinical PVs. A tumor-aware regression model is finally trained by projecting pre-clinical and human tumor transcriptomics data on this consensus representation. **(C)** In order to validate our model, we use positive controls from independent data sources such as copy number or mutation data. These positive controls are established biomarker-drug associations. We compare the predictions of our model to predictions obtained based on these independent established biomarkers. Red boxes highlight our contributions.

This work contains the following novel contributions. First, we introduce a scalable and flexible methodology to find the common factors between pre-clinical models and human tumors. Second, we use this methodology to quantify the transcriptional commonality in biological processes between cell lines, PDXs and human tumors, and we show that these common factors are biologically relevant. Third, we show how these common factors can be used in regression pipelines to predict drug response in human tumors and that we recover well-known biomarker-drug associations. Finally, we derive an equivalent, faster and more interpretable way to compute the geodesic flow kernel, a widely used domain adaptation method in computer vision.

## 2 Material and Methods

### 2.1 Notes on transcriptomics data

We here present the datasets employed in this study. Further notes on pre-processing can be found in Subsection Supp1.3.

#### 2.1.1 The cell line dataset

We used the GDSC1000 dataset to train predictors on the cell lines. GDSC1000 contains IC_50_-values for a wide range of drugs. Amongst these drugs, we restricted ourselves to drugs that are either cytotoxic chemotherapies or targeted therapies and have shown an effect on at least one cancer type. This resulted in a set of 45 drugs employed in this study (Subsection Supp1.2). The gene expression profiles of 1,031 cell lines are available in total, including 51 breast cancer and 40 skin melanoma lines. Gene expression data are available in the form of FPKM.

#### 2.1.2 The PDX dataset

We used the Novartis PDXE dataset which includes the gene expression profiles of 399 PDXs, including 42 breast cancer PDXs and 32 skin melanoma PDXs. Transcriptomics data are available in the form of FPKM.

#### 2.1.3 The human tumor dataset

We extracted gene expression profiles for human tumors from TCGA. Specifically, we employed gene expression profiles for 1,222 breast cancers and 472 skin melanoma cancers. Both FPKM and read counts are available. Mutation and copy number aberrations have been downloaded from the cBioPortal (Gao *et al.* [2013]). Translocation data has been downloaded from TumorFusions (Hu *et al.* [2017]).

### 2.2 The cosine similarity matrix

Transcriptomics data are high-dimensional with *p* ~ 19.000 features (genes) and since these genes are highly correlated, only some combinations of genes are informative. A simple – yet robust (Van Der Maaten *et al.* [2009]) – way to find these combinations is to use a linear dimensionality reduction method, like PCA, that breaks the data matrix down in *d*_*f*_ factors independently for the source (cell lines or PDXs) and target (human tumors) such that

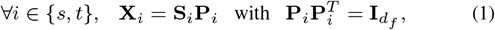

where *s* and *t* refer to the source and target, respectively; **X** represents the (*n* x *p*) transcriptomics dataset where each row represents a sample and each column a gene; 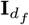 is the identity matrix and 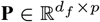 contains the *factors* in the rows (i.e. the principal components). Since these factors are computed independently for the source and the target, we refer to them as *domain-specific factors*. Here we only consider PCA (Principal Components Analysis) since it is widely adopted by the community, and for its direct link to variance that acts as a first-order approximation in the comparison of distributions. Our method is, however, flexible and any linear dimensionality reduction method can be used.

Once domain-specific factors have been independently computed for both the source and target, a simple way to map the source factors to the target factors is to use the *subspace alignment* approach suggested by (Fernando *et al.* [2013]). This approach finds a linear combination (**M**^*^) of source factors that reconstructs the target factors as closely as possible:

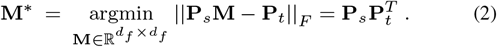

This optimal transformation consists of the scalar product between the source and target factors and therefore quantifies the similarity between the factors. We will therefore refer to it as the *cosine similarity matrix*. It is also referred to as Bregman matrix divergence in the literature.

### 2.3 Common signal extraction by transformation analysis

As we will show in Subsection 3.1, matrix **M**^*^ is far from diagonal, indicating that there is not a one-to-one correspondence between the source-and target-specific factors. Moreover, using **M**^*^ to map the source-projected data onto the target domain-specific factors would only remove source-specific variation, leaving target-specific factors and the associated variation untouched.

To understand this transformation further, we performed a SVD, i.e. 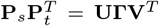, where **U** and **V** are orthogonal of size *d*_*f*_ and **Γ** is a diagonal matrix. **U** and **V** define orthogonal transformations on the source and target domain-specific factors, respectively, and create a new basis for the source and target domain-specific factors:

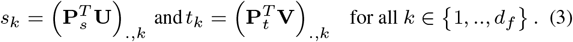

These define the *principal vectors* (PVs) (Golub and Van Loan [2012]) that have the following equivalent definition:

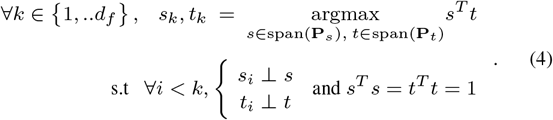

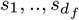 define the same span as the source-specific factors – and so do 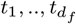 with the target-specific factors. PVs thus retain the same information as the original domain-specific factors, but their cosine similarity matrix (**Γ**) is diagonal. The PVs 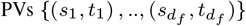 are derived from the source and target domain-specific factors and the pairs are sorted in decreasing order based on their similarity. The top PVs are very similar between source and target while the bottom pairs are very dissimilar. For this reason we restricted the analysis to the top *d*_*pv*_ principal vectors. In (Equation 4), PVs have been defined as unitary vectors that maximise the inner product. The similarities therefore range between 0 and 1, and can thus be interpreted as the cosines of *principal angles* defined as

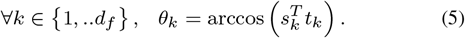

We define **Q**_*s*_ and **Q**_**t**_ as the matrix with the ordered principal vectors of the source and the target, respectively, with the factors in the rows.

### 2.4 Factor-level Gene Set Enrichment Analysis

In order to associate the principal vectors and the consensus representation with biological processes, we use Gene Set Enrichment Analysis (Subramanian *et al.* [2005]). For each factor (i.e. a principal vector or a consensus factor), we projected the tumor data onto it, yielding one score per tumor sample. These sample scores where then used in the GSEA package as continuous phenotypes. We employed sample-level permutation to assess significance based on 1000 permutations. We used two curated gene sets from the MSigDB package: the canonical pathways (cp) and the chemical and genetic perturbations (cgp).

#### Algorithm 1 PRECISE

~~~
**Require:** source data **X**_*s*_, target data **X**_*t*_, number of *domain-specific factors d*_*f*_, number of *principal vector d*_*pv*_.
 **P**_*s*_ ← *d*_*f*_ source *domain-specific factors* (e.g. Principal Components)
 **P**_*t*_ ← *d*_*f*_ target *domain-specific factors* (e.g. Principal Components)
 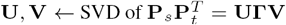
 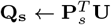
 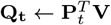
 **for** *i* ← 1 to *d*_*pv*_ **do**
  **S**_*i*_ ← [Φ_*i*_ (0), Φ_*i*_ (0.01), .., Φ_*i*_ (1)]^*T*^
  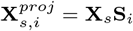
  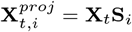
  *τ*_*i*_ ← time of optimal matching between columns of 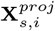 and 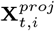
**end for**
**F** ← [Φ_1_ (*τ*_1_), Φ_2_ (*τ*_2_), .., Φ_*d*_ (*τ*_*d*_)]
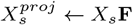
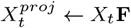
Train a regression model on 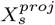
Apply it on the projected target data 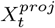
~~~

### 2.5 Building a robust regression model

Given the common factors, we can create a drug response predictor based on these pairs of PVs. There are different ways to use these pairs of PVs. We could restrict ourselves to either the source or target PVs, but it would only support one of the two domains. Alternatively, we could use both source and target PVs. However, this would also be sub-optimal for the following reason. Source PVs are computed using the source data and maximize the explained variance of the source. Hence the source data projected on the source PVs is likely to have higher variance than the source data projected on the target PVs, since target PVs have not been optimized for the source data. If we were to apply penalized regression on the source data projected on both the source and target PVs, it would preferably select the source PVs. This would in turn lead to a loss of generalizability as source-specific information is weighed more heavily that target specific information.

One way to circumvent this issue is to construct a new feature space using ‘intermediate’ features based on interpolation between the spaces spanned by the the source and target PVs. For instance, in the plane that joins *s*_1_ and *t*_1_, the rotations from the former to the latter vector could contain a better representation. The intermediate features are expected to be domain invariant as they represent a trade-off between source and target domains and can thus be used in a regression model.

There is an infinite number of parameterizations for the intermediate features that join the source to the target PVs. As suggested in (Gopalan *et al.* [2011], Gong *et al.* [2012]), we consider the geodesic flow representing the shortest path on the Grassmannian manifold. We derive (see Subsection Supp2.2 for the complete proof) a parameterization of the geodesic as a function of the PVs. Let’s define **Π** and **Ξ** as

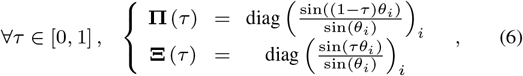

where diag (·) is the diagonal matrix. The intermediate representations are then defined by the geodesic path, that corresponds to a rotation for each pair of PVs:

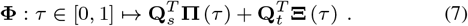

This geodesic path contains, for each pair of PVs, the features forming a rotating arc between the source and the target PVs. This formulation of the geodesic flow has the advantage of being based on the PVs, and not the domain-specific factors, in contrast to the formulation used in (Gopalan *et al.* [2011]) and (Gong *et al.* [2012]). Non-similar PVs can be removed prior to interpolation.

However, we show in Subsection Supp2.3 and Subsection Supp2.4 that projecting on all these features is equivalent, even in the infinite case, to projecting onto both the source and the target principal vectors, with the undesirable consequences described above. It is therefore preferable to create, for each pair of PVs, a single interpolated feature that strikes the right balance between the information contained in the source and target spaces. Consequently, a regression model trained on source data projected on the interpolated feature would generalize better on the target space. To construct these interpolated, or *consensus features*, we use the Kolmogorov-Smirnov (KS) statistic as a measure of similarity between the source and target data, both projected on a candidate interpolated feature.

Specifically, for *i* ∈ {1, *.., d*} and *τ* ∈ [0, 1] the feature at position *τ* between the *i*^*th*^ pair is defined as Φ_*i*_ (*τ*) = (**Φ** (*τ*))_*i*_. For each pair of PVs, we then select the position *τ*_*i*_ that minimizes the KS statistic between the distributions of the source and target data projected on this feature. Let’s denote by *D* the KS statistics between the two projected datasets. We thus define *τ*_*i*_ as:

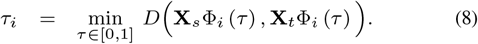

This optimization is performed using a uniformly spaced grid search in interval [0, 1] with step size 0.01, moving between the source and the target.

This process is repeated for each of the top *d*_*pv*_ PVs, resulting in an optimal interpolation position for each. These positions are then plugged back into the geodesic curve to yield the domain-invariant feature representation **F** defined as:

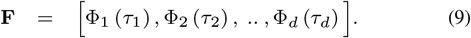

The source data can now be projected on these features and the resulting data set can be used for training a regression model that can be more reliably transferred to the target (human tumor) data.

### 2.6 Notes on implementation

Once the number of principal components and principal vectors have been set (see Subsection Supp5 for an example), the only hyper-parameter that needs to be optimized is the shrinkage coefficient (λ) in the regression model. We employed a nested 10-fold cross validation for this purpose. Specifically, for each of the outer cross validation folds, we employed an inner 10-fold cross-validation on 90% of the data (the outer training fold) to estimate the optimal λ. To this end, in each of the 10 inner folds, we estimated the common subspace, projected the inner training and test fold on the subspace, trained a predictor on the projected inner training fold and determined the performance on the projected inner test fold as a function of λ. After completing these steps for all 10 inner folds, we determined the optimal λ across these results. Then we trained a model with the optimal λ on the outer training fold and applied the predictor to the remaining 10% of the data (outer test fold). We then employed the Pearson correlation between the predicted and actual values on the outer test folds as a metric of predictive performance. Note that every sample in an outer test folds is never employed to perform either domain adaptation nor in constructing the response predictor in that same fold.

The methodology presented in this section is available as a Python 3.7 package available on our GitHub page. The domain adaptation step has been fully coded by ourselves and the regression and cross-validation uses scikit-learn 0.19.2 (Pedregosa *et al.* [2011]).

## 3 Results

### 3.1 Pre-clinical models and human tumors show limited similarity

The cosine similarity matrix **M**^*^ presented in Subsection 2.2 gives an indication of the similarity between the source and the target principal components. A clear correspondence between factors would yield a diagonal matrix, allowing a single target principal component to be assigned to a single source principal component. Using data presented in Subsection 2.1, Fig 2A and Fig 2C instead, show that this is clearly not the case as each source principal component shows similarity to a number of target principal components. Roughly speaking, for cell lines (Fig 2A), the top four source factors show high similarity with the top ten target factors. The similarity between PDXs and human tumor principal components is generally higher and holds for a larger set of factors (Fig 2C). This is to be expected since PDXs are believed to show higher resemblance to human tumors than cell lines.

**Fig. 2.**
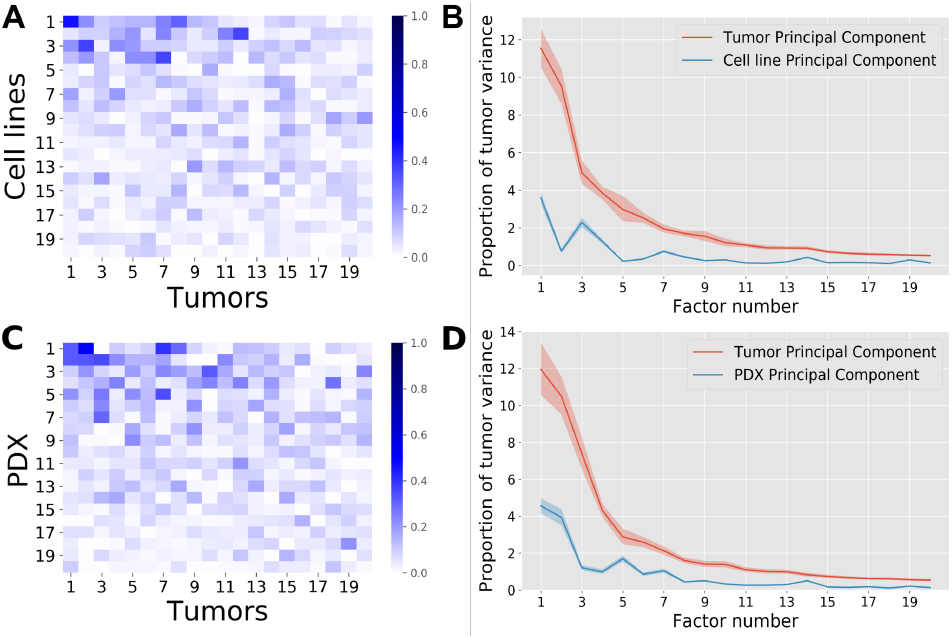
Comparison of domain-specific factors between source and target with the source being cell lines **(A,B)** or PDXs **(C,D)**. **(A,C)** The absolute cosines similarity, i.e. the absolute values of the scalar products between source and target PVs. High similarities are found between some factors but no clear 1-1 correspondence is visible (absence of high values on the diagonal). PDXs show slightly higher similarity to human tumors than cell lines. **(B,D)** The ratio of tumor variance explained. Human tumor data were projected on the source and target PVs, the variance of the projected data on each direction was computed and divided by the total human tumor variance. The shaded regions represent the 98% confidence intervals obtained by bootstrapping the human tumor samples. Overall, the first five cell line factors each explain more than 1% of the human tumor variance each, while for PDXs this is achieved for the first seven factors. The non-monotonic behavior of the PDX and cell line principal component curves shows that the human tumor variance is not supported by the same directions than the pre-clinical models, which necessitates domain adaptation.

When tumor data is projected on the cell line principal components, the explained variance accounts for around 30% of the variance explained when mapping the data on the human tumor principal components (Fig 2B). For PDXs, on the other hand, this amounts to 40% (Fig 2D), again indicating that PDXs resemble tumors more closely than cell lines. The bootstrap confidence intervals obtained by bootstrapping the tumor samples show that the obtained variance proportions differ significantly (Fig 2B and Fig 2D). However, for both model systems, the explained variance is relatively small, indicating that the data for both model systems are not drawn from the same probability distribution as the human tumors, underscoring the need for a proper alignment of the datasets prior to transferring a predictor from the pre-clinical modes to the human tumors.

In order to show that some gene-level structure is shared between these systems, we permuted the order of the genes in the source data only. We then computed the cosine similarities and target explained variance as before (Subsection Supp3.2). Neither the cosine similarity values, nor the variance explained were as high as for the original unshuffled data, suggesting that model systems and tumor cells do share some feature-level structure. We also compared the target data to samples drawn uniformly from a gaussian with a random covariance matrix in order to study whether the similarity between source and target data is significant. As shown in (Subsection Supp3.3), this also yields values three to four orders of magnitude lower than observed. This all shows the existence of a shared signal between source and target.

### 3.2 Principal vectors capture common biological processes

The source and target principal components are employed to compute the ‘common factors’ or PVs for both the source and target (see Subsection 2.3). Fig 3A shows that the source and target PVs exhibit a perfect one-to-one correspondence. This is not unexpected since, by construction, this cosine similarity matrix is the central diagonal matrix (**Γ**) in the SVD of the optimal transformation between source and target (Equation 2). The source and target PVs are directions that respectively support the source and the target variance and are ranked by their pairwise similarity. When 20 principal components are computed between breast cell lines and breast tumors, the similarity between the principal vectors ranges from 0.78 to 0.02, indicating that the low ranking pairs are almost orthogonal. Although 0.78 could seem like a low value for the top similarity, such a value remains significantly large for a 19, 000-dimensional space. Fig Supp5A depicts the top 20 PVs obtained for breast PDXs and breast tumors and shows that the top similarity coefficients are higher than for cell lines, as expected.

**Fig. 3.**
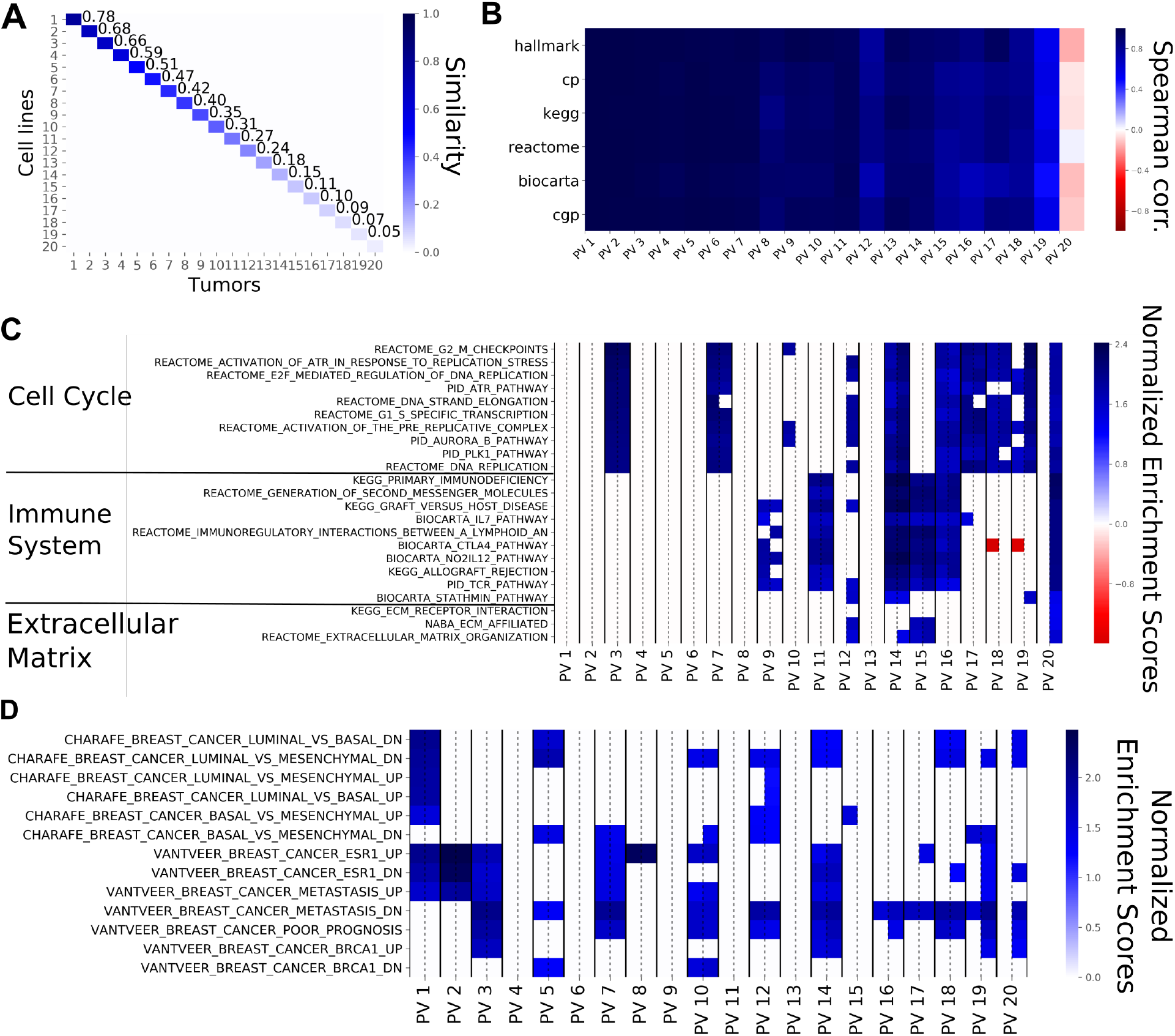
Principal vectors (PVs) computed from breast cell lines and breast tumors from 20 principal components. **(A)** The Cosine Similarity matrix for cell line and tumor PVs. The values on the diagonal show the similarities within the corresponding pairs of PVs. Similarity starts at 78% and goes down to 2% for the last pair (not shown). The off-diagonal values are almost zero, showing that pairs of PVs of unequal rank are orthogonal to one another. **(B)** The Pearson correlations between the Normalized Enrichment Scores (NES) of source and target PVs for the different gene sets employed. The top principal vectors show similar enrichments while the bottom ones show little similarity, even negative correlation. This shows that top principal vectors represent the same biological phenomena. **(C)** The NES based on the Canonical Pathways for each PV pair with the NES for the source PV on the left and the NES for the target PV on the right (separated by a dashed line). Non-significant gene sets are represented as white cells. For this figure panel, we selected the ten gene sets that were most highly enriched in the first five PVs, the ten gene sets that showed the highest enrichment in the bottom PVs as well as all the gene sets related to extra-cellular matrix. The top PVs are exclusively enriched in pathways related to cell cycle. Immune system-related pathways are enriched in the middle and bottom PVs and PVs at the bottom tend to show enrichment for the target PVs only. **(D)** The NES for each PV as displayed in **(C)**, for the CHARAFE and VANTVEER gene sets. The top principal vectors are significantly enriched in sets associated with breast cancer subtypes.

To determine how PVs are related to genes, we calculated the contribution of each gene to the PVs. We subsequently employed these contributions in a gene set enrichment analysis (see Subsection 2.4) to compute the association between the pathways in a given data base and the PVs. This resulted in a vector of pathway scores for every PV. We then computed the pathway similarity between a pair of PVs as the correlation of the pathway scores. Fig 3B shows these correlations for the top 20 pairs of cell line and human tumor PVs for six different pathway databases. Correlations are close to one for the first principal vectors, indicating high pathway similarities, whereas the pathway similarity decreases (even becoming negative) for the lower ranked PVs. Taken together this shows that the principal vectors capture shared pathway information between the model systems and the human tumors.

When zooming in on the individual pathway similarities, we observe roughly two types of behaviors (Fig 3C and Fig 3D). First, some gene sets show significant enrichment for the top PVs as well as lower ranked PVs. These gene sets are related to breast cancer subtypes, cell cycle and DNA replication, i.e. gene sets one would expect to be enriched in both cell lines and human tumors. Second, some gene sets show enrichment for the more dissimilar PVs. Most of these gene sets are related to the response of the immune system and the extra-cellular matrix, entities which are not fully present in the cell lines. Fig Supp5C shows the results of the gene set enrichment analysis for breast PDXs and human tumors. We observe roughly the same behavior as for the cell lines and human tumors, especially regarding the gene sets enriched in the top PVs and the enrichment of immune related sets. However, we do observe that the extra-cellular matrix shows enrichment in higher ranked PVs (PV 5 and 7) which is in line with what one would expect in a PDX model. Taken together, this indicates that the PVs that are most similar between pre-clinical models and human tumors provide a mechanism to capture the information shared between model systems are tumors, while discarding processes that behave differently.

### 3.3 The consensus representation yields reduced but competitive performance

Using PRECISE, we have derived a consensus feature representation which is both biologically informative and shared between the source and target. This representation can be used in a regression model trained on the source drug response data. Since (Jang *et al.* [2014]), demonstrated that regularized linear models such as Ridge regression or ElasticNet (Zou and Hastie [2005]) yield state-of-the-art performance for drug response prediction and since it is widely used, we will be employing Ridge regression.

We computed the predictive performance of PRECISE for 84 drug and tumor type combinations (Subsection Supp1.2). For a given drug and tumor type combination, we used PRECISE with all cell lines (across all tumor types) as source and the corresponding tumor type as target. For example, to predict response to Vemurafenib in melanoma, we used all cell lines, regardless of tumor type, as source, and melanoma tumors from the TCGA as target. We used 70 principal components since the variance explained shows a plateau after 70 principal components (Subsection Supp5.1). For the selection of the top PVs, we compared the obtained similarity values of the source and target PVs to the similarities obtained with random data and put the cutoff at the top 40 PVs (Subsection Supp5.2). We then computed the interpolated feature space employing the KS statistic and employed this feature space in the subsequent Ridge regression models.

As shown in Fig 4A, PRECISE achieves predictive performances that are reduced but comparable with a Ridge regression model trained on the raw cell line gene expression data. The Pearson correlation between Ridge regression and PRECISE performance is *r* = 0.97 with a median relative reduction in Pearson correlation of 0.039. As it is the aim of the consensus representation to focus on the commonalities between the cell lines and the human tumors, it is to be expected that such a representation will not fully capture the variation in the cell lines as it is also adapted to the variation in the human tumors. Hence, a small drop in performance is not unexpected, as long as it results in improved predictions on the human tumor data, which we will demonstrate in the next section.

**Fig. 4.**
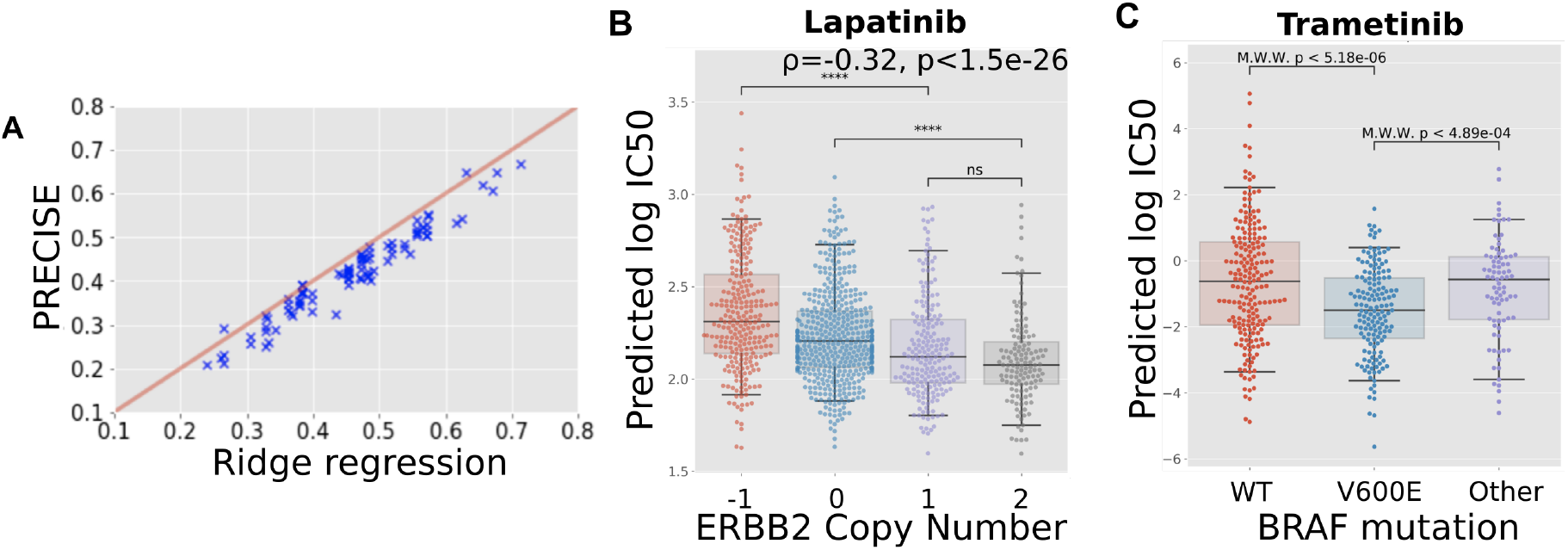
Predictive performance assessment. **(A)** Scatterplot of the performance of PRECISE and Ridge regression. Predictive performance for each approach was computed as the Pearson correlation between the measured and the 10-fold cross-validated predicted IC_50_ s. PRECISE and Ridge regression performance is strongly correlated, with PRECISE showing a slight drop in performance. **(B)** Predicted drug response (predicted IC_50_) for breast tumors based on a predictor of Lapatinib response trained on all the cell lines employing gene expression data only. ERBB2 copy number status in tumors correlates significantly with predicted IC_50_ values, validating the predictions, as tumors with ERBB2 amplifications are known to be more sensitive to Lapatinib. **(C)** Predicted drug response (predicted IC_50_) for skin melanoma tumors based on a predictor of Tramatinib response trained on all the cell lines employing gene expression data only. The PRECISE predictions are validated by the established fact that BRAF^V600E^-mutated tumors show a significantly higher sensitivity while the tumors bearing other mutations in BRAF do not show responses that differ from wild-type tumors.

### 3.4 Domain-invariant regression models recover biomarker-drug associations

To validate our predictor on human tumor data, we follow (Geeleher *et al.* [2017]) by comparing the prediction in human tumors with the performance of independent, known biomarkers. Since we exclusively use gene expression to stratify patients in terms of their response to a certain therapy, we can use known biomarkers derived from other data sources – e.g. mutations or copy number changes – to create an independent response stratification of the human tumors against which we can compare our approach.

The first known and clinically employed association between a biomarker and response to a drug that we tested is the association between the presence of an ERBB2 amplification and response to Lapatinib in breast cancer. Since Lapatinib specifically targets the ERBB2 growth factor, breast tumors that overexpress this growth factor, and are therefore addicted to this signal, respond well to this therapy. An accurate stratification of the breast tumors by PRECISE would therefore show a negative association with the level of ERBB2 amplification in the tumors, as the tumors predicted to be most sensitive (lowest IC_50_) would show the highest level of ERBB2 copy number amplification. This is exactly what we observe in Fig 4B, where the predicted IC_50_ and the observed ERBB2 copy number amplification level show a Spearman correlation of *ρ* = −0.32. In addition, the copy number loss and copy number neutral categories show statistically significant differences in predicted IC_50_ compared to the samples harboring a copy number gain.

The second known association that we investigated is the association between the presence of a BRAF^V600E^ mutation and response to MEK inhibitors in skin melanomas, while other BRAF mutations have not exhibited this association. As shown in Fig 4C, the IC_50_s predicted for Trametinib by PRECISE show a significant difference between tumors bearing a BRAF^V600E^ mutation and tumors that are wild type for this gene. In addition, the PRECISE predictions also show a significant difference in predicted Trametinib response between tumors bearing a BRAF^V600E^ mutation and tumors bearing other mutations in this gene.

Other known associations we tested against are shown in Fig Supp9: Dabrafenib sensitivity predicted by BRAF^V600E^ mutations(Fig Supp9A), Vemurafenib sensitivity predicted by BRAF^V600E^ (Fig Supp9B) mutations, Imatinib sensitivity predicted by BCR/ABL translocations in Acute Myeloid Leukemia (Fig Supp9C), Olaparib sensitivity predicted by BRCA1 deletion in breast cancer (Fig Supp9D) and Talazoparib sensitivity predicted by BRCA1 deletion in breast cancer (Fig Supp9E). These results have been computed using all the cell lines as source. When using solely cell lines from the tissue under consideration, the same results are observed, with a larger difference in predicted sensitivity for BRAF^V600E^ mutated tumors for Trametinib (Fig Supp9G), Dabrafenib (Fig Supp9H) and Vemurafenib (Fig Supp9I), and a very clear separation of BCR/ABL translocated tumors from the rest under Imatinib treatment (Fig Supp9J).

Finally, we replicated results from (Geeleher *et al.* [2017]) by employing Ridge regression on either the raw or ComBat corrected gene expression data. Lapatinib sensitivity is predicted as well as by PRECISE (Fig Supp10A) with or without the ComBat pre-processing step. For Dabrafenib, the association with BRAF^V600E^ is recovered, with and without ComBat. However, while the strength of the association obtained with ComBat is comparable to the association recovered with PRECISE, the strength diminishes without ComBat pre-processing (Fig Supp10C). In contrast to PRECISE, neither Ridge regression nor Ridge regression in combination with ComBat were able to recover the association between the BRAF^V600E^ mutation and Trametinib.

In summary, PRECISE is capable of retrieving all tested associations between known biomarkers and drug response, while current state-of-the-art approaches fail to recover all these associations.

## 4 Discussion

Using high throughput sequencing and screening technologies, scientists have leveraged the versatility of pre-clinical models over the past decade to create powerful predictors of drug response. However, due to the intrinsic differences between cell lines, PDXs and real human tumors, these predictors can not be expected to directly translate to the human setting. We have quantified the overlap in terms of the transcriptomics signal between these pre-clinical models and human tumors. We then introduced PRECISE, a domain adaptation framework that finds shared mechanisms between pre-clinical models and human tumors that display the same behavior across these systems.

PRECISE generates PVs, pairs of factors that capture the common variance between pre-clinical models and human tumors. The top pairs of PVs are most similar, and thus recapitulate molecular behavior shared between the systems. The least similar PVs, can be discarded since they correspond to mechanisms not shared across systems.

These vectors depend on the choice of linear dimensionality reduction methods employed. We employed PCA, but other methods have been proposed in the literature (e.g. Bismeijer *et al.* [2018], Argelaguet *et al.* [2018]), all having different qualities (e.g. being biologically meaningful, filtering out noise, etc.). The versatility of our method enables the use of any dimensionality reduction scheme, as long it finds an informative linear subspace.

Interpolation between the source and target principal vectors gives rise to features that balance the contribution of pre-clinical models and tumors. We showed (Section Supp2) that interpolating between the principal vectors is equivalent to employing the Geodesic Flow Kernel approach that relies on the geodesics on the Grassmannian manifold (Gong *et al.* [2012]) and has already yielded state-of-the-art performance in Computer Vision.

Recently, other ways to interpolate between the source and the target domains have been proposed, such as (Caseiro *et al.* [2015]) that use a spline instead of the geodesic. We devised a simple, yet effective interpolation scheme between the source and target PVs where we employed the similarity based on the Kolmogorov-Smirnov statistic between the source and target data projected on the interpolated space to arrive at a consensus representation. This representation strikes the right balance between the pre-clinical models and the human tumors.

We subsequently projected the data on this consensus representation and trained a regression model which takes the distribution of the tumor gene expression data into account. This work considered Ridge and ElasticNet, but our approach is versatile and can be employed in combination with any classification or regression approach.

We showed that a Ridge regression model based on the consensus representation achieves slightly reduced performance compared to state-of-the-art approaches applied directly to the raw cell line gene expression data. This is to be expected, as the consensus representation filters out cell line specific information while capturing more relevant tumor variation, hence enabling efficient transfer to the tumor samples.

We finally compared our predictions to the performance of known biomarkers such as BRAF^V600E^ in skin cancer or ERBB2 amplification in breast cancer and show that our method can reliably recover the associations between these biomarkers and their companion drugs. We show that response to Lapatinib can be predicted better when all cell lines are used for domain adaptation. On the other hand, using all cell lines reduced the power to predict response to Vemurafenib, although the resulting association with BRAF^V600E^ mutation status remained significant. This might be due to the ubiquity of the ERBB2 amplification in several tumor types, in contrast to the BRAF^V600E^ mutation that is specific to particular tumor types.

We restricted our study to domain adaptation based on transcriptomics data only, as it has been shown to be the most predictive data type (Jang *et al.* [2014]). However, dissimilar behavior between pre-clinical models and human tumors might also be present in other molecular data types. A multi-omic drug response predictor should also correct for these differences, which will require a multi-omic domain adaptation approach which accommodates the unique data characteristics of each molecular data type.

Other methods have recently been proposed to tackle the problem of transferring pre-clinical predictors to human tumors. In (Webber *et al.* [2018]), the authors create a correlation network for each omics data type and jointly map these networks onto a protein-protein interaction network. They then select the cliques that are conserved across omics-layers. In (Normand *et al.* [2018]), the authors present an elegant framework for fold-change prediction in humans based on data from mouse models. Using fold-change data from both humans and mouse, a linear model is fitted at the gene-level. This linear model is then used to predict fold changes in human tumors for novel conditions. The problem of translating from model systems to human has broad applications and we envision that it will be a very active area of future research.

## Acknowledgements

We thank Mirrelijn van Nee, Gergana Bounova, Daniel Vis, Stavros Makrodimitris, Tycho Bismeijer and Mark van de Wiel for useful discussions. We thank Nanne Aben, Marie Corradie and Wouter Kouw for critical reading of the manuscript and useful feedback on the methodology.

## Footnotes

1 While this terminology is not widely used in the community, we follow the categorization employed in (Pan and Yang [2010]).

2 Our method, although using the notion of canonical angle, differs significantly from the Canonical Correlation Analysis. Indeed, in our case, samples are not paired and no cross-correlation can be computed.

## References

Argelaguet, R., Velten, B., Arnol, D., Dietrich, S., Zenz, T., Marioni, J. C., Buettner, F., Huber, W., and Stegle, O. (2018). Multi-omics factor analysis—a framework for unsupervised integration of multi-omics data sets. Molecular systems biology, 14(6), e8124.

Ben-David, U., Ha, G., Tseng, Y.-Y., Greenwald, N. F., Oh, C., Shih, J., McFarland, J. M., Wong, B., Boehm, J. S., Beroukhim, R., et al. (2017). Patient-derived xenografts undergo mouse-specific tumor evolution. Nature genetics, 49(11), 1567.

Ben-David, U., Siranosian, B., Ha, G., Tang, H., Oren, Y., Hinohara, K., Strathdee, C. A., Dempster, J., Lyons, N. J., Burns, R., et al. (2018). Genetic and transcriptional evolution alters cancer cell line drug response. Nature, page 1.

Bismeijer, T., Canisius, S., and Wessels, L. F. (2018). Molecular characterization of breast and lung tumors by integration of multiple data types with functional sparse-factor analysis. PLoS computational biology, 14(10), e1006520.

Caseiro, R., Henriques, J. F., Martins, P., and Batista, J. (2015). Beyond the shortest path: Unsupervised domain adaptation by sampling subspaces along the spline flow. In Proceedings of the IEEE Conference on Computer Vision and Pattern Recognition, pages 3846–3854.

Csurka, G. (2017). Domain adaptation for visual applications: A comprehensive survey. arXiv preprint arXiv:1702.05374.

Duan, L., Tsang, I. W., and Xu, D. (2012). Domain transfer multiple kernel learning. IEEE Transactions on Pattern Analysis and Machine Intelligence, 34(3), 465–479.

Fernando, B., Habrard, A., Sebban, M., and Tuytelaars, T. (2013). Unsupervised visual domain adaptation using subspace alignment. In Proceedings of the IEEE international conference on computer vision, pages 2960–2967.

Gao, H., Korn, J. M., Ferretti, S., Monahan, J. E., Wang, Y., Singh, M., Zhang, C., Schnell, C., Yang, G., Zhang, Y., et al. (2015). High-throughput screening using patient-derived tumor xenografts to predict clinical trial drug response. Nature medicine, 21(11), 1318.

Gao, J., Aksoy, B. A., Dogrusoz, U., Dresdner, G., Gross, B., Sumer, S. O., Sun, Y., Jacobsen, A., Sinha, R., Larsson, E., et al. (2013). Integrative analysis of complex cancer genomics and clinical profiles using the cbioportal. Sci. Signal., 6(269), p11–p11.

Geeleher, P., Cox, N. J., and Huang, R. S. (2014). Clinical drug response can be predicted using baseline gene expression levels and in vitro drug sensitivity in cell lines. Genome Biology.

Geeleher, P., Zhang, Z., Wang, F., Gruener, R. F., Nath, A., Morrison, G., Bhutra, S., Grossman, R. L., and Huang, R. S. (2017). Discovering novel pharmacogenomic biomarkers by imputing drug response in cancer patients from large genomics studies. Genome Research.

Gillet, J.-P., Varma, S., and Gottesman, M. M. (2013). The Clinical Relevance of Cancer Cell Lines. JNCI Journal of the National Cancer Institute.

Golub, G. H. and Van Loan, C. F. (2012). Matrix computations, volume 3. JHU Press.

Gong, B., Shi, Y., Sha, F., and Grauman, K. (2012). Geodesic flow kernel for unsupervised domain adaptation. In Computer Vision and Pattern Recognition (CVPR), 2012 IEEE Conference on, pages 2066–2073. IEEE.

Gopalan, R., Li, R., and Chellappa, R. (2011). Domain adaptation for object recognition: An unsupervised approach. In Computer Vision (ICCV), 2011 IEEE International Conference on, pages 999–1006. IEEE.

Hu, X., Wang, Q., Tang, M., Barthel, F., Amin, S., Yoshihara, K., Lang, F. M., Martinez-Ledesma, E., Lee, S. H., Zheng, S., et al. (2017). Tumorfusions: an integrative resource for cancer-associated transcript fusions. Nucleic acids research, 46(D1), D1144–D1149.

Iorio, F., Knijnenburg, T. A., Vis, D. J., Bignell, G. R., Menden, M. P., Schubert, M., Aben, N., Gonçalves, E., Barthorpe, S., Lightfoot, H., Cokelaer, T., Greninger, P., van Dyk, E., Chang, H., de Silva, H., Heyn, H., Deng, X., Egan, R. K., Liu, Q., Mironenko, T., Mitropoulos, X., Richardson, L., Wang, J., Zhang, T., Moran, S., Sayols, S., Soleimani, M., Tamborero, D., Lopez-Bigas, N., Ross-Macdonald, P., Esteller, M., Gray, N. S., Haber, D. A., Stratton, M. R., Benes, C. H., Wessels, L. F., Saez-Rodriguez, J., McDermott, U., and Garnett, M. J. (2016). A Landscape of Pharmacogenomic Interactions in Cancer. Cell.

Jang, I. S., Neto, E. C., Guinney, J., Friend, S. H., and Margolin, A. A. (2014). Systematic assessment of analytical methods for drug sensitivity prediction from cancer cell line data. In Biocomputing 2014, pages 63–74. World Scientific.

Normand, R., Du, W., Briller, M., Gaujoux, R., Starosvetsky, E., Ziv-Kenet, A., Shalev-Malul, G., Tibshirani, R. J., and Shen-Orr, S. S. (2018). Found in translation: a machine learning model for mouse-to-human inference. Nature methods, 15(12), 1067.

Pan, S. J. and Yang, Q. (2010). A survey on transfer learning. IEEE Transactions on knowledge and data engineering, 22(10), 1345–1359.

Pan, S. J., Kwok, J. T., and Yang, Q. (2008). Transfer learning via dimensionality reduction. In AAAI, volume 8, pages 677–682.

Pan, S. J., Tsang, I. W., Kwok, J. T., and Yang, Q. (2011). Domain adaptation via transfer component analysis. IEEE Transactions on Neural Networks, 22(2), 199–210.

Pedregosa, F., Varoquaux, G., Gramfort, A., Michel, V., Thirion, B., Grisel, O., Blondel, M., Prettenhofer, P., Weiss, R., Dubourg, V., Vanderplas, J., Passos, A., Cournapeau, D., Brucher, M., Perrot, M., and Duchesnay, E. (2011). Scikit-learn: Machine learning in Python. Journal of Machine Learning Research, 12, 2825–2830.

Song, L., Gretton, A., Borgwardt, K. M., and Smola, A. J. (2008). Colored maximum variance unfolding. In Advances in neural information processing systems, pages 1385–1392.

Subramanian, A., Tamayo, P., Mootha, V. K., Mukherjee, S., Ebert, B. L., Gillette, M. A., Paulovich, A., Pomeroy, S. L., Golub, T. R., Lander, E. S., et al. (2005). Gene set enrichment analysis: a knowledge-based approach for interpreting genome-wide expression profiles. Proceedings of the National Academy of Sciences, 102(43), 15545–15550.

Van Der Maaten, L., Postma, E., and Van den Herik, J. (2009). Dimensionality reduction: a comparative review. Journal of Machine Learning Research, 10, 66–71.

Webber, J. T., Kaushik, S., and Bandyopadhyay, S. (2018). Integration of tumor genomic data with cell lines using multi-dimensional network modules improves cancer pharmacogenomics. Cell systems, 7(5), 526–536.

Zou, H. and Hastie, T. (2005). Regularization and variable selection via the elastic net. Journal of the Royal Statistical Society: Series B (Statistical Methodology), 67(2), 301–320.

