## Supplementary_Text for "PRECISE: A domain adaptation approach to transfer predictors of drug response from pre-clinical models to tumors"

#### Supplementary Material

Soufiane Mourragui, Marco Loog, Marcel JT Reinders, Lodewyk FA Wessels

#### Supp1 Additional information

##### Supp1.1 Notations

| Notation | Meaning |
|---|---|
| $\mathbb{R}$ | Real numbers |
| $.^T$ | Transposition operation |
| $\|\| \cdot \|\|_F$ | Frobenius norm |
| $\mathbb{G}(d, p)$ | Grassmann manifold of parameters $d$ and $p$ (for $d < p$) |
| span | Span of a set of vectors |
| $()_{\cdot,k}$ | $k^{\text{th}}$ column of the matrix |
| argmax | Element that maximises the following quantity |
| argmin | Element that minimizes the following quantity |
| arcos | Inverse trigonometric function of the cosine |
| $p$ | Number of genes ($\sim 19,000$) |
| $\cdot_s$ | Related to the source dataset (cell line or PDX) |
| $\cdot_t$ | Related to the target dataset (tumor) |
| $n_\cdot$ | Number of samples |
| $d_f$ | Number of domain-specific factors, e.g. principal components |
| $d_{pv}$ | Number of principal vectors |
| $\mathbf{X}_\cdot \in \mathbb{R}^{n_\cdot p}$ | Transcriptomics data with samples in the rows |
| $\mathbf{I}_d$ | Identity matrix of size $d \times d$ |
| diag | Diagonal matrix |
| $\mathbf{P}_\cdot \in \mathbb{R}^{d_f p}$ | Domain-specific factors with factors in the rows. |
| $\mathbf{M}^*$ | Cosine similarity matrix |
| $\theta$ | Principal angle |
| $\mathbf{Q}_\cdot \in \mathbb{R}^{d_{pv} p}$ | Principal vectors with factors in the rows |
| $\mathbf{\Phi}$ | Geodesic in Grassmann manifold |
| $\mathbf{\Pi}$ | Source-related rotation for geodesic in Grassmannian |
| $\mathbf{\Xi}$ | Target-related rotation for geodesic in Grassmannian |
| $\tau \in [0, 1]$ | Interpolation time between source and target |
| $\Phi_i,\ i \in \{1, .., d_{pv}\}$ | Interpolation between the $i^{\text{th}}$ pairs of principal vectors |
| $D$ | Kolmogorov-Smirnov (KS) statistics |
| $\tau_i$ | Time of optimal matching for $i^{\text{th}}$ pairs of principal vectors |
| $\mathbf{F}$ | Consensus representation |
| $\mathbf{X}_\cdot^{proj}$ | Data projected on the consensus representation $\mathbf{F}$. |

#### Supp1.2 List of drugs

| Drug | Cancer types used in PRECISE |
|---|---|
| Erlotinib | Head and Neck |
| Sunitinib | Colorectal |
| Paclitaxel | Breast, Ovary, Lung |
| Cyclopamine | Breast |
| AX628 | Lung |
| Sorafenib | Kidney |
| Crizotinib | Lung |
| S-Trityl-L-cysteine | Prostate |
| Parthenolide | Blood |
| Roscovitine | Lung |
| Salubrinal | Lung |
| Lapatinib | Breast |
| Doxorubicin | Breast, Bladder |
| Etoposide | Testis, Lung, Brain, Ovary |
| Gemcitabine | Breast, Ovary, Lung, Pancreas, Bladder |
| Mitomycin C | Oesophagus |
| Vinorelbine | Breast, Lung |
| Bicalutamide | Prostate |
| 5-Fluorouracil | Colorectal, Oesophagus, Stomach, Skin |
| Thapsigargin | Brain |
| Bleomycin | Testis, Ovary, Cervix |
| Pazopanib | Kidney |
| Zibotentan | Prostate |
| Camptothecin | Ovary, Lung |
| Vinblastine | Lung, Bladder, Brain, Skin, Testis |
| Cisplatin | Testis, Ovary, Cervix, Breast, Bladder, Head and Neck, Brain, Oesophagus |
| Docetaxel | Breast, Head and Neck, Stomach, Prostate, Lung |
| Methotrexate | Breast |
| Gefitinib | Breast, Lung |
| Nilotinib | Blood |
| RDEA119 | Skin |
| Temsirolimus | Kidney |
| Olaparib | Ovary, Breast, Prostate |
| Lenalidomide | Blood |
| Axitinib (+ rescreen) | Kidney |
| Elesclomol | Skin |
| Afatinib | Lung |
| Vismodegib | Skin |
| Cetuximab | Colorectal, Lung, Head and Neck |
| Tamoxifen | Breast |
| Trametinib | Skin |
| Dabrafenib | Skin |
| Temozolomide | Brain |
| AZD6244 | Lung |

#### Supp1.3 Notes on transcriptomics data

Transcriptome levels have been measured using RNA-Seq Illumina HTSeq for both cell lines, PDX as well as the tumors. For cell lines and tumors, RNA-Seq data was available as read counts. For PDX and tumors, RNA-Seq data was available as FPKM. Since FPKM values are corrected for gene length at the transcript level and already normalised for library size, they cannot directly be compared to read counts.

Consequently, we use two separate pre-processing pipelines, following the recommendation in (Dillies *et al.* (2013); Zwiener *et al.* (2014)). For read counts, data is first normalized using TMM (Robinson and Oshlack (2010)), then log-transformed and mean-centered. For FPKM, data is log-transformed and mean-centered. Experiments involving cell line to human tumor transfer have been performed using read counts, while PDX to human tumor transfer experiments have been performed using FPKM.

#### Supp2 Geodesic Flow derivation

##### Supp2.1 Original formulation

We denote by $\mathbb{G}(d,p)$ the Grassmannian of $d$-dimensional subspaces within a $p$-dimensional space. This is formally defined as the set with a Riemannian structure of the $d$-dimensional subspaces within a larger $p$-dimensional space. The geometry of this space is non-Euclidean and therefore the shortest paths to go from one point to another are referred to as geodesic. The source *domain-specific factors* can be represented by one point in the Grassmannian and so do the target *domain-specific factors*. The idea now is to find this geodesic within $\mathbb{G}(d,p)$ that links the two. An analytical formulation of this curve is given in (Gong *et al.* (2012)).
A SVD on the cosine similarity matrix yields the matrices $\mathbf{U}_1 \in \mathbb{R}^{d\times d}$ and $\mathbf{U}_2 \in \mathbb{R}^{(p-d)\times d}$ such that

$$\mathbf{P}_s^T\mathbf{P_t} = \mathbf{U}_1\mathbf{\Gamma}\mathbf{V}^T \quad \text{where} \quad \mathbf{\Gamma} = \text{diag}\left(\cos\left(\theta_1\right),..,\cos\left(\theta_d\right)\right) \tag{Supp1}$$

Let $\mathbf{R}_s \in \mathbf{R}^{p\times(p-d)}$ be the orthonormal complement of $\mathbf{P}_s$ (i.e. $\mathbf{P}_s^T\mathbf{R}_s = \mathbf{0}_{p-d,d}$ and $\mathbf{R}_s^T\mathbf{R_s} = \mathbf{I}_{p-d,p-d}$). The cosine similarity matrix between the orthogonal complement of $\mathbf{P}_s$ and the matrix $\mathbf{P}_t$ gives, after a SVD and a column-wise permutation on the right matrix:

$$\mathbf{R}_s^T\mathbf{P_t} = -\mathbf{U}_2\mathbf{\Sigma}\mathbf{V}^T \quad \text{where} \quad \mathbf{\Sigma} = \text{diag}\left(\sin\left(\theta_1\right),..,\sin\left(\theta_d\right)\right) \tag{Supp2}$$

With these quantities, one can now define:

**Proposition Supp2.1** (Geodesic on the Grassmann manifold)**.** *The geodesic on the Grassmann manifold can be represented by the bases $\mathbf{\Phi}$ defined as:*

$$\begin{aligned} \mathbf{\Phi}: \quad & [0,1] \longrightarrow \mathbb{G}(d,p) \\ & \tau \longmapsto \mathbf{P}_s\mathbf{U}_1\mathbf{\Gamma}(\tau) - \mathbf{R}_s\mathbf{U}_2\mathbf{\Sigma}(\tau) \\ \textit{where} \quad & \mathbf{\Gamma}(\tau) = \textit{diag}\left(\cos\left(\tau\theta_1\right),..,\cos\left(\tau\theta_d\right)\right) \\ \textit{and} \quad & \mathbf{\Sigma}(\tau) = \textit{diag}\left(\sin\left(\tau\theta_1\right),..,\sin\left(\tau\theta_d\right)\right) \end{aligned} \tag{Supp3}$$

As shown in (Equation Supp3), this formulation requires a lot of computation since the orthogonal complement $\mathbf{R}_s$ has to be computed. What is more, it links the *domain-specific factors* together, which is of limited interest for our study. Indeed, we would like to have a formulation that directly links the *principal vectors* instead, in order to filter out irrelevant factors that are too dissimilar to be used in the regression model.

##### Supp2.2 Writing the geodesic flow in terms of principal vectors instead of principal components

We here derive a formulation of the geodesic $\mathbf{\Phi}$ in terms of principal vectors. We only make the assumption that $\theta_d < \frac{\pi}{2}$, which can easily be checked experimentally, which generally holds for all practical purposes. For problems that nevertheless do not satisfy this assumption, orthogonal principal vectors can be removed from the problem. They indeed do not correspond to transferable features and can be discarded.

**Proposition Supp2.2** (Equivalent definition of the Geodesic)**.** *Let's assume that $\theta_d < \frac{\pi}{2}$, then the geodesic can equivalently be defined as*

$$\begin{aligned} \forall\tau \in [0,1], \quad \mathbf{\Phi}(\tau) &= \mathbf{Q}_s\mathbf{\Pi}(\tau) + \mathbf{Q}_t\mathbf{\Xi}(\tau) \\ \textit{with} \quad \mathbf{\Pi}(\tau) &= \textit{diag}\left(\frac{\sin\left(\left(1-\tau\right)\theta_i\right)}{\sin\left(\theta_i\right)}\right) \\ \textit{and} \quad \mathbf{\Xi}(\tau) &= \textit{diag}\left(\frac{\sin\left(\tau\theta_i\right)}{\sin\left(\theta_i\right)}\right) \end{aligned} \quad . \tag{Supp4}$$

*Proof.* Since $[\mathbf{P}_s, \mathbf{R}_s]$ forms a orthogonal basis of $\mathbb{R}^p$, we have $\mathbf{P}_s\mathbf{P}_s^T + \mathbf{R}_s\mathbf{R}_s^T = \mathbf{I}_p$. Summing up then (Equation Supp1) and (Equation Supp2) yields, after multiplying by $\mathbf{P}_s^T$ and $\mathbf{R}_s^T$ respectively:

$$\mathbf{P}_t\mathbf{V} = \mathbf{P}_s\mathbf{U}_1\mathbf{\Gamma} - \mathbf{R}_s\mathbf{U}_2\mathbf{\Sigma} \tag{Supp5}$$

We find that $\mathbf{\Phi}(1) = \mathbf{P}_t\mathbf{V} = \mathbf{Q}_t$, which means that the end point of the geodesic gives us the basis of target principal vectors. Since $\theta_d < \frac{\pi}{2}$, then $\forall i \in \{1,..,d\}, \theta_i < \frac{\pi}{2}$. $\mathbf{\Sigma}$ will thus be invertible and (Equation Supp5) yields:

$$-\mathbf{R}_s\mathbf{U}_2 \quad = \quad \mathbf{Q}_t\mathbf{\Sigma}^{-1} - \mathbf{Q}_s\mathbf{\Gamma}\mathbf{\Sigma}^{-1} \tag{Supp6}$$

Plugging (Equation Supp6) into (Supp3) yields the desired formula. ■

This way, the geodesic path is computed in $\mathcal{O}(p \times d)$ instead of $\mathcal{O}(p^2)$ and does not require the computation of the orthogonal complement – which can be computationally intensive. This formulation has the interest of taking the principal vectors as inputs, instead of the principal components. It shows that the geodesic interpolates between principal vectors within each pair by taking features forming a rotating arc between the source and the target principal vectors. It therefore proves that our approach using all the principal vectors is strictly similar to the approach proposed in (Gong *et al.* (2012)) and in (Gopalan *et al.* (2011)).

#### Supp2.3 Equivalence between Geodesic Flow Sampling and Principal Vector regression

As suggested by (Gopalan *et al.* (2011)), a domain-invariant drug response predictor can be created by sampling the interval $[0,1]$, i.e. by taking a number $M+1$ of intermediate representations $\left\{0, \frac{1}{M}..., 1\right\}$, computing the corresponding intermediate features $\left\{\mathbf{\Phi}(0), \mathbf{\Phi}\left(\frac{1}{M}\right)..., \mathbf{\Phi}(1)\right\}$, and finally projecting source and target data on these intermediate features. We show here that it strictly equivalent to projecting on the principal vectors and learn a linear regression model onto these principal vectors.

**Proposition Supp2.3** (Equivalence of estimators without penalization)**.** *Let $\hat{y}_S$ be the linear drug response estimator learnt without penalization by minimizing the loss function $\ell$ on the interpolated coefficients, and let $\hat{y}_{PV}$ be the linear estimator learnt by minimizing the loss function $\ell$ on the principal vectors. Then, $\hat{y}_S = \hat{y}_{PV}$.*

*Proof.* Let $x \in \mathbb{R}^p$ be a sample - from either source or target. A linear model learnt on the projected data will give a response of the form:

$$\begin{aligned}
& \hat{y}_S\left(x; (\alpha_{i,j})_{\substack{1\le i\le d\\0\le j\le M}}\right) \\
= & \sum_{i=1}^{d}\sum_{j=0}^{M}\alpha_{i,j}x^T\left(\mathbf{Q}_{s,i}\mathbf{\Pi}_{i,i}\left(\frac{j}{m}\right) + \mathbf{Q}_{t,i}\mathbf{\Xi}_{i,i}\left(\frac{j}{m}\right)\right) \\
= & \sum_{i=1}^{d}x^T\left[\mathbf{Q}_{s,i}\sum_{j=0}^{M}\alpha_{i,j}\mathbf{\Pi}_{i,i}\left(\frac{j}{m}\right) \quad +\mathbf{Q}_{t,i}\sum_{j=0}^{M}\alpha_{i,j}\mathbf{\Xi}_{i,i}\left(\frac{j}{m}\right)\right] \\
= & \sum_{i=1}^{d}x^T\left[\beta_i^s\mathbf{Q}_{s,i} + \beta_i^t\mathbf{Q}_{t,i}\right] \\
= & \hat{y}_{PV}\left(x; \left(\beta_i^s, \beta_i^t\right)_{1\le i\le d}\right)
\end{aligned} \tag{Supp7}$$

with:

- $\alpha_{i,j} \in \mathbb{R}$ for all $i \in \{1,..,d\}$ and $j \in \{0,..,M\}$ the coefficients of the linear model fitted on the interpolated features.
- $\forall i \in \{1,..,d\}, \beta_i^s = \sum_{j=0}^{M}\alpha_{i,j}\mathbf{\Pi}_{i,i}\left(\frac{j}{m}\right)$
- $\forall i \in \{1,..,d\}, \beta_i^s = \sum_{j=0}^{M}\alpha_{i,j}\mathbf{\Xi}_{i,i}\left(\frac{j}{m}\right)$

Therefore, using this reciprocal correspondence, we can state that the non-regularized minimization procedure, using any loss $\ell$ is equivalent for both set of parameters, namely:

$$\min_{\alpha_{i,j}}\frac{1}{n}\sum_{k=1}^{n}\ell\big(y_k,\ \hat{y}_S(x_k;\alpha_{i,j})\big) \ = \ \min_{\beta_i^s,\beta_i^t}\frac{1}{n}\sum_{k=1}^{n}\ell\big(y_k,\ \hat{y}_{PV}\left(x_k;\beta_i^s,\beta_i^t\right)\big) \tag{Supp8}$$

■

Penalization may change the matter and the solution of the two minimization procedure might change slightly. However, we advocate for the latter penalized minimization procedure. Indeed, only $2d$ parameters have to be penalized. This in turn makes the minimization procedure easier and numerically more stable. The former formulation would require shrinking on way more features that are expressing the same content (same total rank).

#### Supp2.4 Equivalent formulation of Geodesic Flow Kernel Matrix

The original definition of the Geodesic Flow Kernel is (Gong *et al.* (2012)):

$$\begin{aligned} \forall x, y \in \mathbb{R}^p, \quad & \int_0^1 \left(\mathbf{\Phi}(\tau)^T x\right)^T \left(\mathbf{\Phi}(\tau)^T y\right) d\tau \; = \; x^T \mathbf{G} y \\ \text{with} \quad \mathbf{G} \; = \; & \begin{bmatrix} \mathbf{P}_s \mathbf{U}_1 & \mathbf{R}_s \mathbf{U}_2 \end{bmatrix} \begin{bmatrix} \mathbf{\Lambda}_1 & \mathbf{\Lambda}_2 \\ \mathbf{\Lambda}_2 & \mathbf{\Lambda}_3 \end{bmatrix} \begin{bmatrix} \mathbf{U}_1^T \mathbf{P}_s^T \\ \mathbf{U}_2^T \mathbf{R}_s^T \end{bmatrix} \end{aligned} \tag{Supp9}$$

As shown in (Equation Supp9), computing the matrix $\mathbf{G}$ requires quadratic time in the number of covariates, which can be prohibitive in genomics (when $p \sim 20,000$). We show here how to improve this computation using the new formulation of (Equation Supp4).

**Proposition Supp2.4** (Equivalent definition of Geodesic Flow Kernel)**.** *If $\theta_d < \frac{\pi}{2}$, then there exists $\sigma_1, .., \sigma_d \in \mathbb{R}$ and $\omega_1, .., \omega_d \in \mathbb{R}$ such that*

$$\mathbf{G} \quad = \quad \begin{bmatrix} \widetilde{\mathbf{Q}_s} \\ \widetilde{\mathbf{Q}_t} \end{bmatrix}^T \begin{bmatrix} \widetilde{\mathbf{Q}_s} \\ \widetilde{\mathbf{Q}_t} \end{bmatrix}$$

*with*

$$\widetilde{\mathbf{Q}_s} = \begin{bmatrix} \mathbf{Q}_{s,1}\sigma_1 + \mathbf{Q}_{t,1}\omega_1 \\ .. \\ \mathbf{Q}_{s,d}\sigma_d + \mathbf{Q}_{t,d}\omega_d \end{bmatrix} \quad \textit{and} \quad \widetilde{\mathbf{Q}_t} = \begin{bmatrix} \mathbf{Q}_{s,1}\omega_1 + \mathbf{Q}_{t,1}\sigma_1 \\ .. \\ \mathbf{Q}_{s,d}\omega_d + \mathbf{Q}_{t,d}\sigma_d \end{bmatrix}$$

*Proof.* First, if $x \in \mathbb{R}^p$, we define $x_s = x^T \mathbf{Q_s}$ and $x_t = x^T \mathbf{Q_t}$ as the projection of the point $x$ and the source and target principal vectors. Then, using flow formulation from (Equation Supp4), we get:

$$\begin{aligned} & \int_0^1 x^T \mathbf{\Phi}(\tau) \mathbf{\Phi}(\tau)^T y \; d\tau \\ = & \int_0^1 x^T \left[\mathbf{Q}_s \mathbf{\Pi}(\tau) + \mathbf{Q}_t \mathbf{\Xi}(\tau)\right] \left[\mathbf{\Pi}(\tau) \mathbf{Q}_s^T + \mathbf{\Xi}(\tau) \mathbf{Q}_t^T\right] y \; d\tau \\ = & \quad x_s^T \left[\int_0^1 \mathbf{\Pi}^2(\tau) \; d\tau\right] y_s \\ & + \; x_t^T \left[\int_0^1 \mathbf{\Xi}^2(\tau) \; d\tau\right] y_t \\ & + \; x_s^T \left[\int_0^1 \mathbf{\Pi}(\tau) \mathbf{\Xi}(\tau) \; d\tau\right] y_t \\ & + \; x_t^T \left[\int_0^1 \mathbf{\Xi}(\tau) \mathbf{\Pi}(\tau) \; d\tau\right] y_s \\ = & \begin{bmatrix} x_s^T & x_t^T \end{bmatrix} \begin{bmatrix} \int_0^1 \mathbf{\Pi}^2(\tau) \; d\tau & \int_0^1 \mathbf{\Xi}(\tau) \mathbf{\Pi}(\tau) \; d\tau \\ \int_0^1 \mathbf{\Xi}(\tau) \mathbf{\Pi}(\tau) \; d\tau & \int_0^1 \mathbf{\Xi}^2(\tau) \; d\tau \end{bmatrix} \begin{bmatrix} y_s \\ y_t \end{bmatrix} \end{aligned} \tag{Supp10}$$

With simple trigonometrical identities, we can show that :

$$\int_0^1 \mathbf{\Pi}^2(\tau) \, d\tau \; = \; \int_0^1 \mathbf{\Xi}^2(\tau) \, dt \; = \; \left(\frac{\theta_i - \sin(\theta_i)\cos(\theta_i)}{2\theta_i \sin^2(\theta_i)}\right)_i \tag{Supp11}$$

$$\int_0^1 \mathbf{\Pi}(\tau)\,\mathbf{\Xi}(\tau)\;d\tau \;=\; \left(\frac{\sin(\theta_i) - \theta_i\cos(\theta_i)}{2\theta_i \sin^2(\theta_i)}\right)_i \tag{Supp12}$$

Since the matrix is diagonal, we now have a formulation that only requires $\mathcal{O}(d + dp)$, faster than the $\mathcal{O}(p^2)$ that we had before since with $d = 50$ and $p = 19000$, we get a 8x speed-up.

We can write the matrix $\mathbf{G}$ as a product of the principal vector instead :

$$\begin{aligned} \mathbf{G} &= \begin{bmatrix}\mathbf{Q}_s^T & \mathbf{Q}_t^T\end{bmatrix}\begin{bmatrix}\mathbf{\Lambda} & \boldsymbol{\mu}\\ \boldsymbol{\mu} & \mathbf{\Lambda}\end{bmatrix}\begin{bmatrix}\mathbf{Q}_s\\ \mathbf{Q}_t\end{bmatrix} \\ \mathbf{\Lambda} &= \text{ diag}\left(\frac{\theta_i - \sin(\theta_i)\cos(\theta_i)}{2\theta_i\sin^2(\theta_i)}\right) \\ \boldsymbol{\mu} &= \text{ diag}\left(\frac{\sin(\theta_i) - \theta_i\cos(\theta_i)}{2\theta_i\sin^2(\theta_i)}\right) \end{aligned} \tag{Supp13}$$

Let's denote $(\lambda_1, ..\lambda_d)$ the diagonal coefficients of $\mathbf{\Lambda}$ and $(\mu_1, ..\mu_d)$ the diagonal coefficients of $\boldsymbol{\mu}$. We can now define the coefficients $\sigma_i$ and $\omega_i$ for all $i \in \{1, .., d\}$ as

$$\begin{aligned} \sigma_i &= \frac{1}{2}\left(\sqrt{\lambda_i + \mu_i} + \sqrt{\lambda_i - \mu_i}\right) \\ \omega_i &= \frac{1}{2}\left(\sqrt{\lambda_i + \mu_i} - \sqrt{\lambda_i - \mu_i}\right) \end{aligned} \tag{Supp14}$$

and the matrix $\mathbf{H}$ as :

$$\mathbf{H} \;=\; \begin{bmatrix} \sigma_1 & & & \omega_1 & & \\ & \cdots & & & .. & \\ & & \sigma_d & & & \omega_d \\ \omega_1 & & & \sigma_1 & & \\ & \cdots & & & .. & \\ & & \omega_d & & & \sigma_d \end{bmatrix} \tag{Supp15}$$

$\mathbf{H}$ is positive semi-definite (symmetric with eigenvalues $\sigma_i + \omega_i > 0$ and $\sigma_i - \omega_i > 0$) and respect:

$$\begin{bmatrix}\mathbf{\Lambda} & \boldsymbol{\mu}\\ \boldsymbol{\mu} & \mathbf{\Lambda}\end{bmatrix} \;=\; \mathbf{H}^T\mathbf{H} \tag{Supp16}$$

Plugging this equality in (Equation Supp13), we get:

$$\mathbf{G} \;=\; \begin{bmatrix}\mathbf{Q}_s^T & \mathbf{Q}_t^T\end{bmatrix}\mathbf{H}^T\mathbf{H}\begin{bmatrix}\mathbf{Q}_s\\ \mathbf{Q}_t\end{bmatrix} \tag{Supp17}$$

Let's now define the two following matrices:

$$\widetilde{\mathbf{Q}_s} = \begin{bmatrix}\mathbf{Q}_{s,1}\sigma_1 + \mathbf{Q}_{t,1}\omega_1\\ ..\\ \mathbf{Q}_{s,d}\sigma_d + \mathbf{Q}_{t,d}\omega_d\end{bmatrix} \quad \text{and} \quad \widetilde{\mathbf{Q}_t} = \begin{bmatrix}\mathbf{Q}_{s,1}\omega_1 + \mathbf{Q}_{t,1}\sigma_1\\ ..\\ \mathbf{Q}_{s,d}\omega_d + \mathbf{Q}_{t,d}\sigma_d\end{bmatrix} \tag{Supp18}$$

We finally get:

$$\mathbf{G} \;=\; \begin{bmatrix}\widetilde{\mathbf{Q}_s}\\ \widetilde{\mathbf{Q}_t}\end{bmatrix}^T \begin{bmatrix}\widetilde{\mathbf{Q}_s}\\ \widetilde{\mathbf{Q}_t}\end{bmatrix} \tag{Supp19}$$

■

The geodesic flow kernel is therefore equivalent to projecting on $2d$ vectors that form a basis equivalent to the source and target principal vectors. Using the same idea as in Prop Supp2.4, the ordinary least square estimate will be equivalent to the one obtained using principal vectors.

### Supp3 Comparison of factors between source and target

#### Supp3.1 Comparison results for other tissues

Following experiments from Fig 2, we computed the cosine similarity and the variance explained for other tissues. Results can be found in Fig Supp1.

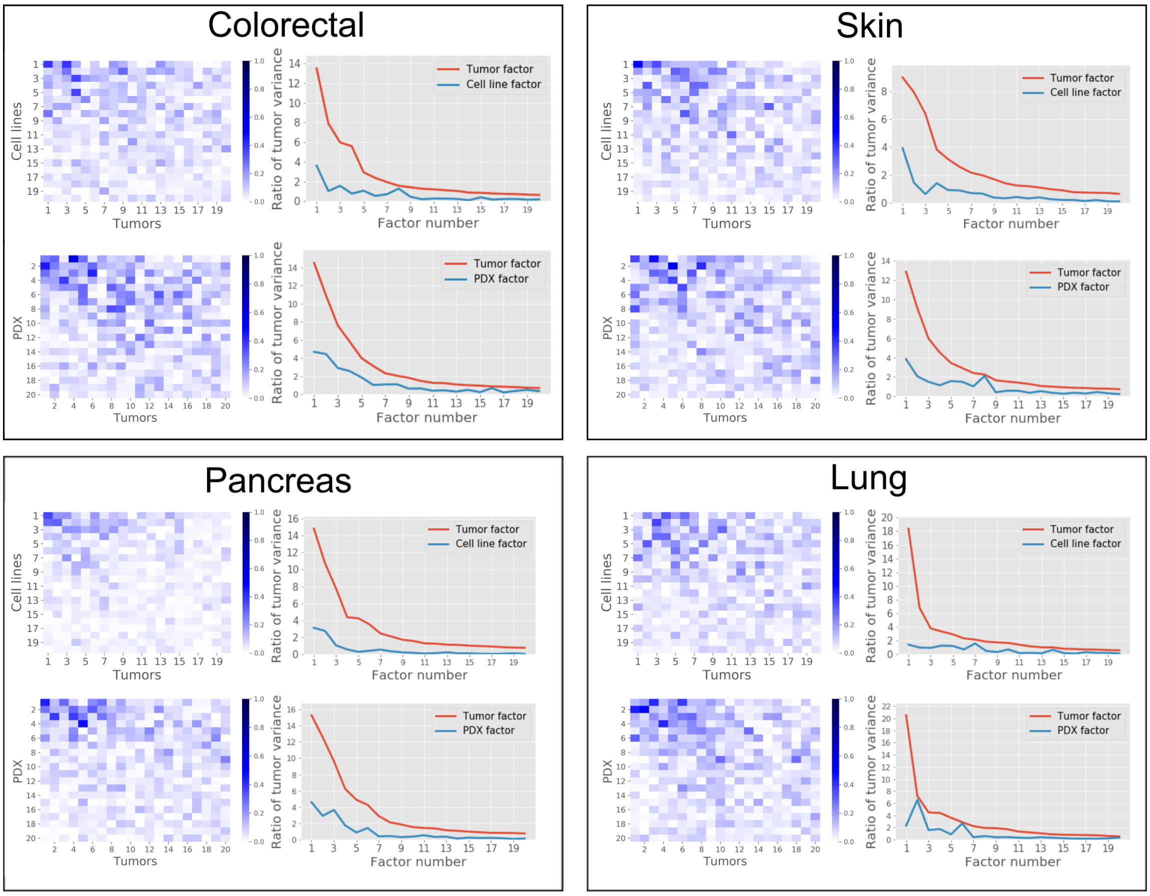


Figure Supp1: **Cosine similarity matrices between pre-clinical models and tumors for different tissues**. Each box represents the analysis carried out on one tissue type. Within each box, the top panel represents the cell line analysis and the bottom one represents the PDX results. Cosine similarity values between source (cell lines or PDXs) and target (tumors) are displayed on the left. Ratio of target variance explained by source principal components is displayed on the right panel.

#### Supp3.2 Significance of the cosine similarity values

To show that these cosine similarity values are significant, we performed a permutation test at the gene level. These cosine similarity values are supposed to reflect some shared structure in the data. If we permute the source genes while keeping the target data intact, this structure should be destroyed. The source principal components would be different and the cosine similarity values should be impacted. We permuted the genes order at the source level only and computed the resulting cosine similarity matrix and variance explained 1000 times to create a meaningful comparison on 5 tissues : breast, colorectal, lung, skin and pancreas. The results are displayed in Fig Supp2.

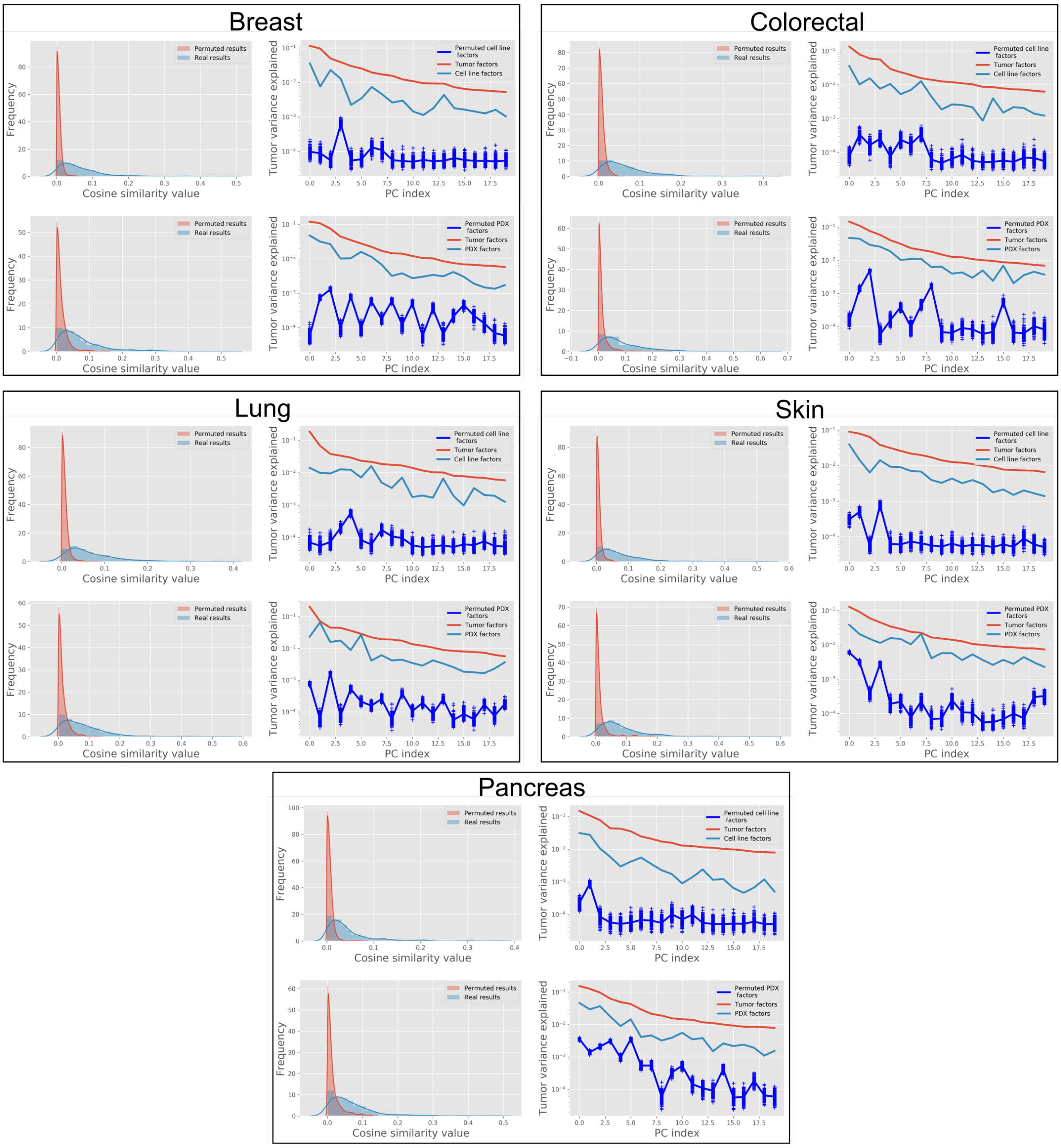


Figure Supp2: **Gene-level permutation results**. For each (source, target) couple, genes have been permuted in the source data. Cosine similarity values and target variance explained have then been computed as in Subsection 3.1. For each tissue, the top row represents results between cell lines and tumors while the bottom one represents results for PDX and tumors. The left column represents the histogram of cosine similarity values while the right column shows the variance explained by target, source and gene-level-permuted source principal components. 1000 permutations have been employed to arrive to these results. For every tissue, the cosine similarity values for the permuted source data range from 0 to 0.05, while certain cosine similarity values are as large as 0.2 for almost every tissue. It suggests that the cosine similarity values encountered in Fig 2 and Fig Supp1 are not the product of non-comparable signals. When it comes to the variance explained, the variance explained by permuted source principal components is consistently two to three orders of magnitude lower than when the tumor data is projected on the non-permuted source data. Two notable exceptions: colorectal PDXs and and Pancreatic PDXs for which some permuted principal components show variance explained only one order of magnitude lower than the non-permuted one.

#### Supp3.3 Comparison with random signals

Gene-level permutation, although yielding useful insights as shown in Subsection Supp3.2, restricts the pool of principal components values to the feature-level permutations. To go one step further in the identification, we used a random signal to quantify the commonality. We computed the cosine similarity values and the tumor variance explained for 250 random covariance matrices using the following protocol:

1. A random covariance matrix was sampled uniformly from the positive semi-definite matrices.
2. 1000 data points were drawn from the Gaussian distribution with 0-mean and the covariance matrix drawn in 1.
3. Principal components were computed from the these data points and cosine similarity values were computed alongside the tumor explained variance and compared to real data.

Although the second step could be removed and principal components could be computed directly using the randomly drawn covariance matrix, we decided to use sampled data to be the closest possible to our original setting. 1000 corresponds to the total number of cell lines available and is therefore comparable to our settings. Results are shown in Fig Supp3.

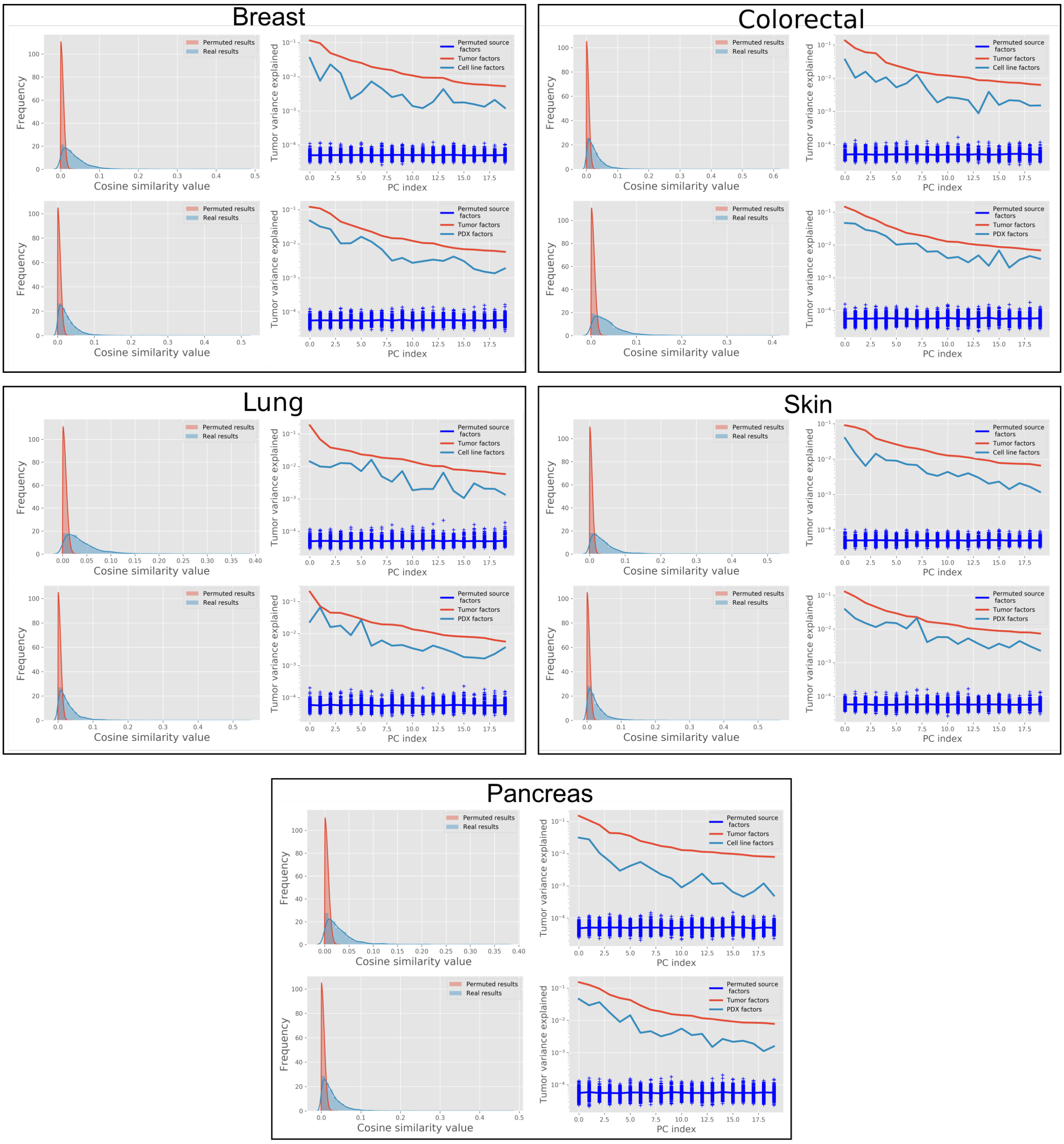


Figure Supp3: **Random signal results**. For each (source, target) couple, 250 positive semi-definite matrices were drawn randomly. For each matrix, 1000 data points were then drawn from a Gaussian distribution with this matrix as covariance. Cosine similarity and tumor variance explained were finally computed. These purely random signals are here compared to the real results. For each tissue, the top panel represents the comparison for cell lines while the bottom represents the results for PDXs. On the left are compared the cosine similarity values and on the right the tumor variance explained ratio. The random cosine similarity values appear to be consistently ranging between 0 and 0.02 while cosine similarity between tumors and real source data are as large as 0.2 for some principal components. It indicates that the similarity values between pre-clinical systems and tumors are not the product of the comparison of two random signals. In terms of variance explained, the variance explained by random principal components is two to five orders of magnitude lower than the tumor variance explained by real source principal components. This result is consistent across all tissue type and once again indicate the existence of some common structure between source and target.

### Supp4 Principal Vectors analysis for different set of tissues

#### Supp4.1 Breast vs Breast for PDX

In Fig 3, we compared breast cancer cell lines to human breast tumors. The same experiment was run using PDXs instead of cell lines and results are shown in Fig Supp5.

#### Supp4.2 Breast vs All

In Fig 4, PRECISE was trained using all cell lines in order to enhance the sample size to around 1000 – only 52 breast cancer cell lines are available. We compared the making of the principal vectors between all cell lines and the breast tumors to make sure that these principal vectors still show some enrichment. Results are shown in Fig Supp5.

#### Supp4.3 Skin vs Skin

We repeated the experiment of Fig 3 to another tissue: skin. As shown in Fig Supp6, the same behavior as in breast appears, with immune related pathways mostly enriched in the least similar PVs.

#### Supp4.4 Other tissue

We computed the similarity scores for other tissues: skin, lung, pancreas and colorectal. Results are shown in Fig Supp7 for both cell lines and PDXs.

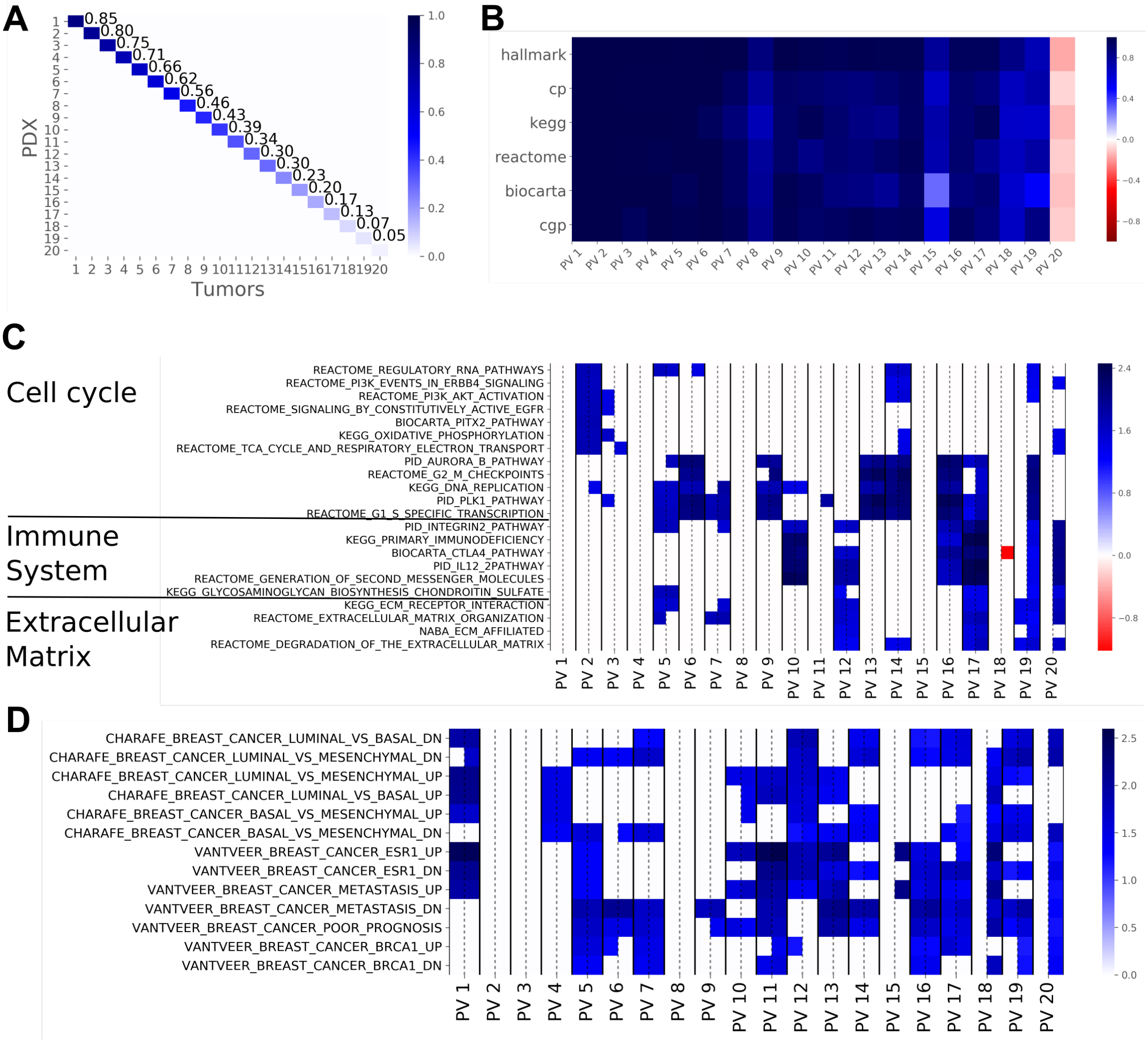


Figure Supp4: **Principal vectors (PVs) computed from breast PDXs and breast tumors from 20 principal components.**. **(A)** Cosine similarity matrix between PDX and tumor principal vectors. As shown on the diagonal, the similarity is higher than in Fig 3A for a similar sample size. This is encouraging since PDXs are expected to mimic human tumors more closely than cell lines. **(B)** Spearman Correlation between PDX and tumor PV Normalized Enrichment Score (NES) for several gene set collections. The spearman correlation is almost 1 up to the 8th PV, suggesting that the same pathways get enriched. The last PV pair shows a negative correlation, in accordance with the almost null similarity. **(C)** The NES based on the Canonical Pathways for each PV pair with the NES for the source PV on the left and the NES for the target PV on the right (separated by a dashed line). Non-significant gene sets are represented as white cells. For this figure panel, we selected the ten gene sets that were most highly enriched in the first five PVs, the ten gene sets that showed the highest enrichment in the bottom PVs as well as all the gene sets related to extra-cellular matrix. The top PVs are exclusively enriched in pathways related to cell cycle. Immune system-related pathways are enriched in the middle and bottom PVs and PVs at the bottom tend to show enrichment for the target PVs only. Compared to results of Fig 2C, the gene sets related to the immune system appear to be again enriched only in the less similar PVs, while extracellular matrix related pathways are this time showing some enrichment for the top PVs. **(D)** The NES for each PV as displayed in **(C)**, for the CHARAFE and VANTVEER gene sets. The top principal vectors are significantly enriched in sets associated with breast cancer subtypes.

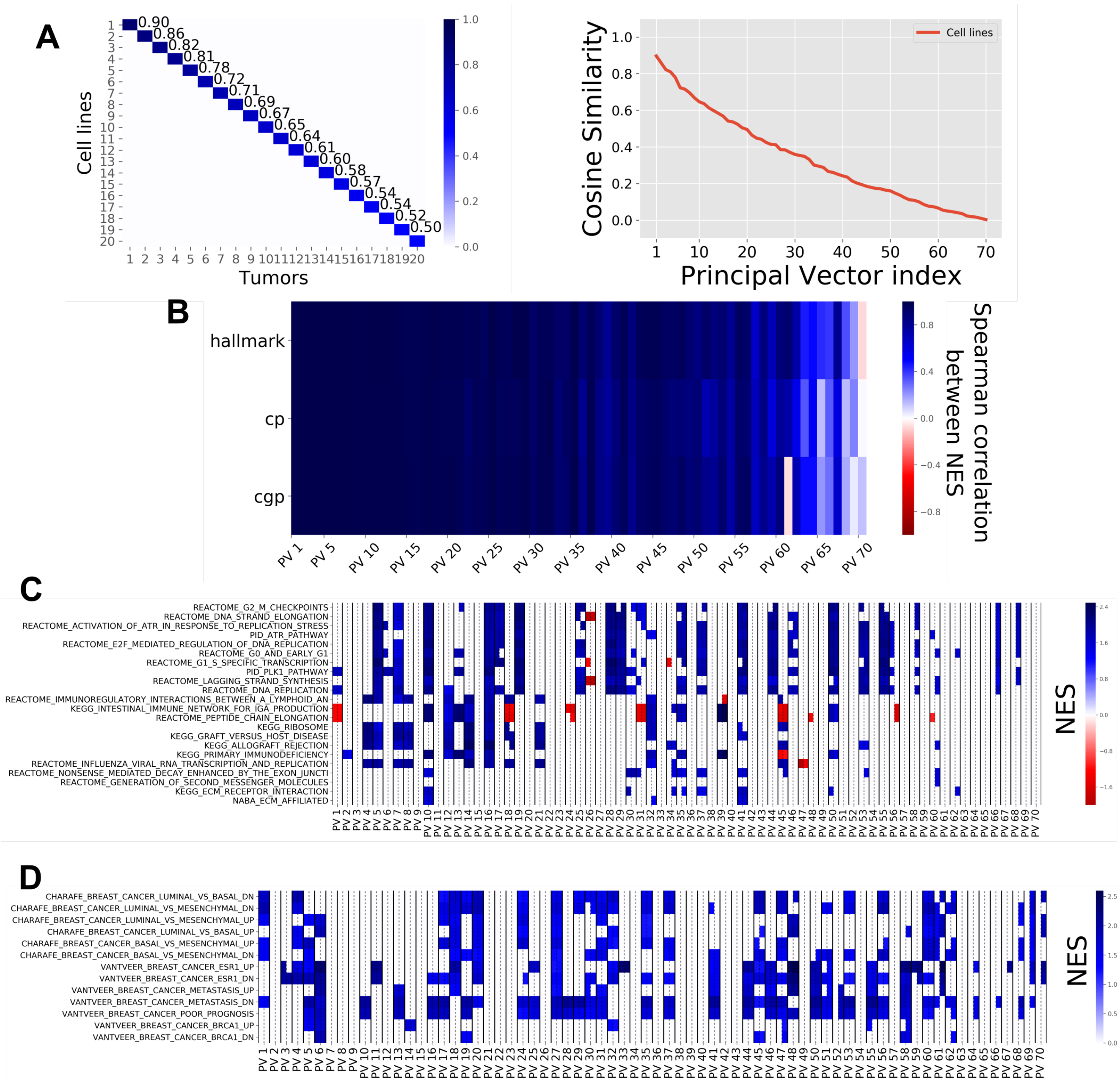


Figure Supp5: **Principal vectors (PVs) computed from breast cell lines and breast tumors from 70 principal components..** **(A)** Cosine similarity values between the top 70 principal vectors. A zoom is performed on the top 20, showing that similarity is as high as 90% for the top pair. **(B)** Spearman correlation between Normalised Enrichment Scores (NES) within each pair of PVs. Correlations close to 1 in the top 30 PV show that gene sets get the same enrichment in cell line and tumor PV and indicate an important structural similarity. **(C)** The NES based on the Canonical Pathways for each PV pair with the NES for the source PV on the left and the NES for the target PV on the right (separated by a dashed line). Non-significant gene sets are represented as white cells. For this figure panel, we selected the ten gene sets that were most highly enriched in the first five PVs, the ten gene sets that showed the highest enrichment in the bottom PVs as well as all the gene sets related to extra-cellular matrix. **(D)** The NES for each PV as displayed in **(C)**, for the CHARAFE and VANTVEER gene sets. The top principal vectors are significantly enriched in sets associated with breast cancer subtypes.

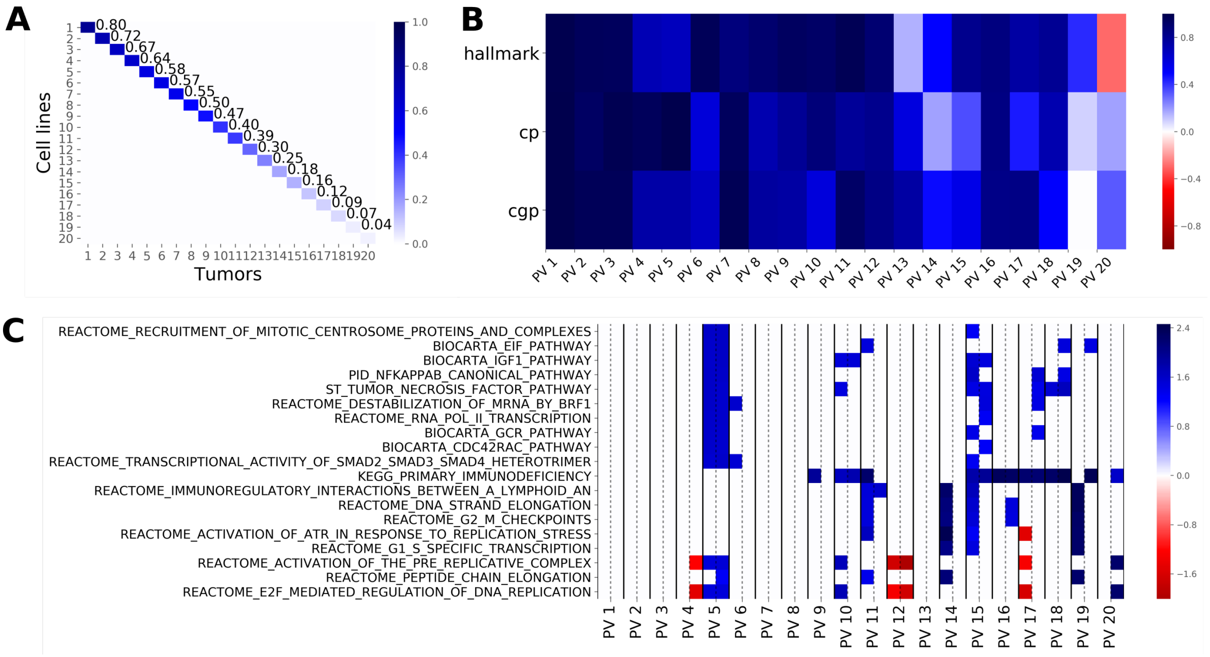


Figure Supp6: **Principal vectors (PVs) computed from skin cell lines and skin tumors from 20 principal components.**. **(A)** Cosine similarity matrix between cell lines and tumor principal vectors. **(B)** Spearman Correlation between PDX and tumor PV Normalized Enrichment Score (NES) for several gene set collections. For the skin, the spearman correlations between NES are lower than for breast, although they remain larger than 0.8 for the top 10 PVs. **(C)** The NES based on the Canonical Pathways for each PV pair with the NES for the source PV on the left and the NES for the target PV on the right (separated by a dashed line). Non-significant gene sets are represented as white cells. For this figure panel, we selected the ten gene sets that were most highly enriched in the first five PVs, the ten gene sets that showed the highest enrichment in the bottom PVs. Although two immune-related pathways are enriched in the top PVs, the same pattern as in Fig 3 appears with several pathways enriched exclusively in the last principal vectors.

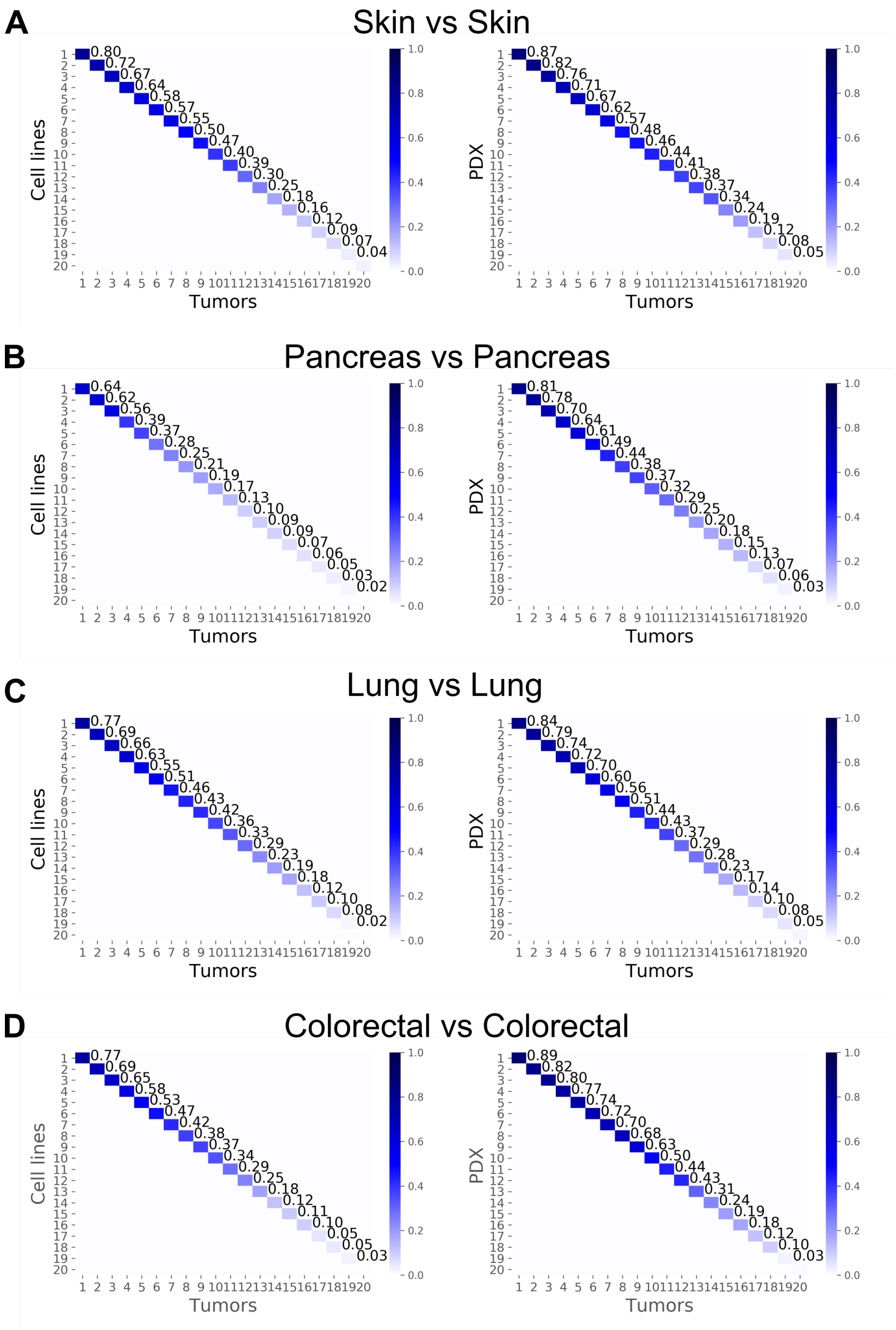


Figure Supp7: **The cosine similarity matrix between principal vectors for other tissues with 20 principal components**. **(A)** Similarity values when target is set as skin tumors and source set as skin cell lines (left) or skin PDXs (right). **(B)** Similarity values when target is set as pancreatic tumors and source set as pancreatic cell lines (left) or pancreatic PDXs (right). **(C)** Similarity values when target is set as lung tumors and source set as lung cell lines (left) or lung PDXs (right). **(D)** Similarity values when target is set as colorectal tumors and source set as colorectal cell lines (left) or colorectal PDXs (right).

#### Supp5 Choice of the hyper parameters for the experiments

##### Supp5.1 Variance-based approach for selecting the number of Principal Components

We selected the number of domain-specific factors (PCs) based on the variance explained by the cell line principal components. Since the sample size is always larger for tumors, this cut-off point is lower for cell lines than for tumors and we only showed the cell line behavior. As shown in Fig Supp8, we took 20 PCs when the same tissue is used for source and for target ; we took 70 PCs when all cell lines are used as source data.

##### Supp5.2 Comparison to the randomly-sampled data for determining the similarity cut-off point

Once the number of PCs had been settled, we needed to determine the number of PVs to select. For that purpose, we computed the similarity between tumor data and data drawn from a gaussian distribution with a random covariance matrix. We repeated this experiment 250 times and got 250 similarity profiles. We took as threshold the maximum random similarity and selected PVs with similarity at least as large. As shown in Fig Supp8, it yields 15 PVs when only one tissue is used for source, and 40 when all cell lines are taken into account.

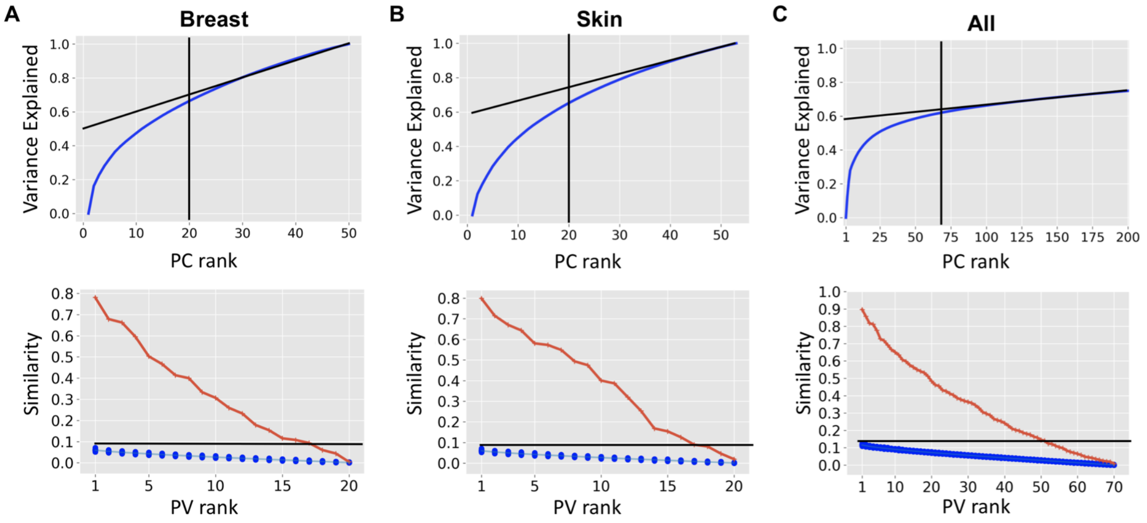


Figure Supp8: **Choice of hyperparameters** $d_f$ **and** $d_{pv}$. The top panel shows the cumulative variance explained, while the bottom panel shows the similarity between the resulting PVs computed from the number of PCs found with the top panel. **(A)** shows results for breast cell lines (with breast tumors), **(B)** shows results for skin cell lines (with skin tumors), and **(C)** shows results for all cell lines (with breast tumors). For selecting the number of PCs, we drew a line corresponding to the asymptotic behavior of the cumulative variance and selected the principal components for which the cumulative variance explained does not follow this behavior. This gives a cut-off slightly before 20 for breast, slightly above 20 for skin and around 70 for all. Since we want to use the same number of PCs for all experiment having one tissue for the source, we settled for 20 that makes consensus between skin and breast. We settled to 70 for experiments with all cell lines. Once this number of PCs had been settled, we needed to determine where to put the PV threshold. For that, we sampled data from random covariance matrix 1000 times and compute 1000 similarity profiles following a similar idea than in Fig Supp3. We take the top similarity as cut-off, which yields 15 PVs for breast, slighlty more for skin and around 40 for PVs. Based on this experiment, we decided to settle for 15 PV when one tissue of the cell line is taken and 40 PVs when all cell lines are taken into account.

### Supp6 Comparison with known biomarkers

#### Supp6.1 PRECISE correlation with other known mechanisms

We repeated the experiment of Fig 4 with other known biomarker-drug associations. We also repeated the same experiments but took only one tissue for the cell lines. Results shown in Fig Supp9 indicate than PRECISE successfully recapitulates known associations coming from independent data sources.

#### Supp6.2 Biomarker correlation for Ridge regression without any domain adaptation or with ComBat as preprocessing step

We compared PRECISE results to the scenario where no domain adaptation is used and a Ridge regression is trained on the cell lines and directly transferred on the human tumors. We also compared PRECISE to the pipeline used in (Geeleher *et al.* (2014)), where the difference between cell lines and human tumors is modelled as a batch effect. As shown in Fig Supp10, most of the associations are still recapitulated by the two scenarios, but PRECISE offers a higher discriminative power on most of the biomarkers.

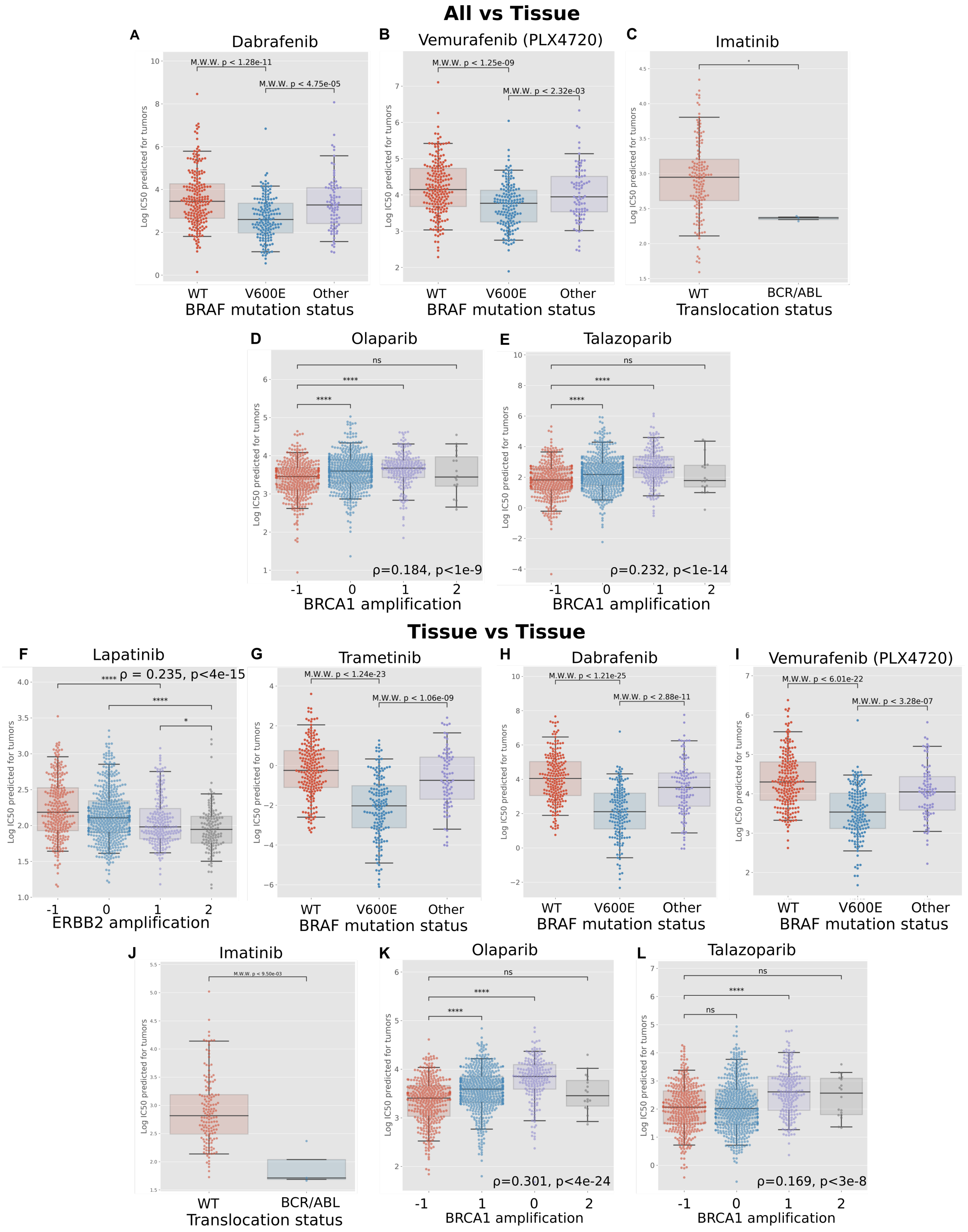


Figure Supp9: **Comparison to other known biomarkers**. Following the same experimental procedure as in Fig 4, we compared $IC_{50}$ predicted by PRECISE with some known biomarkers. Using all cell lines as source data, we show that PRECISE prediction are validated in **(A)** Dabrafenib (sensitive to $BRAF^{V600E}$ mutation), **(B)** Vemurafenib (sensitive to $BRAF^{V600E}$ mutation), **(C)** Imatinib (sensitive to BCR/ABL translocation), **(D)** Olaparib (sensitive to BRCA1 deletion) and **(E)** Talazoparib (sensitive to BRCA1 deletion). We repeated the experiment using only one tissue type in cell lines with all of the investigated drugs. We show that using only breast cell lines reduces the predicted power of ERBB2 in Lapatinib **(F)** and of BRCA1 in Talazoparib **(L)**. However, it increases the power of $BRAF^{V600E}$ mutation in all the MEK inhibitors considered **(G,H,I)**, completely discriminates BCR/ABL translocated tumors for Imatinib **(J)** and increases the power of BRCA1 deletion in Olaparib **(K)**.

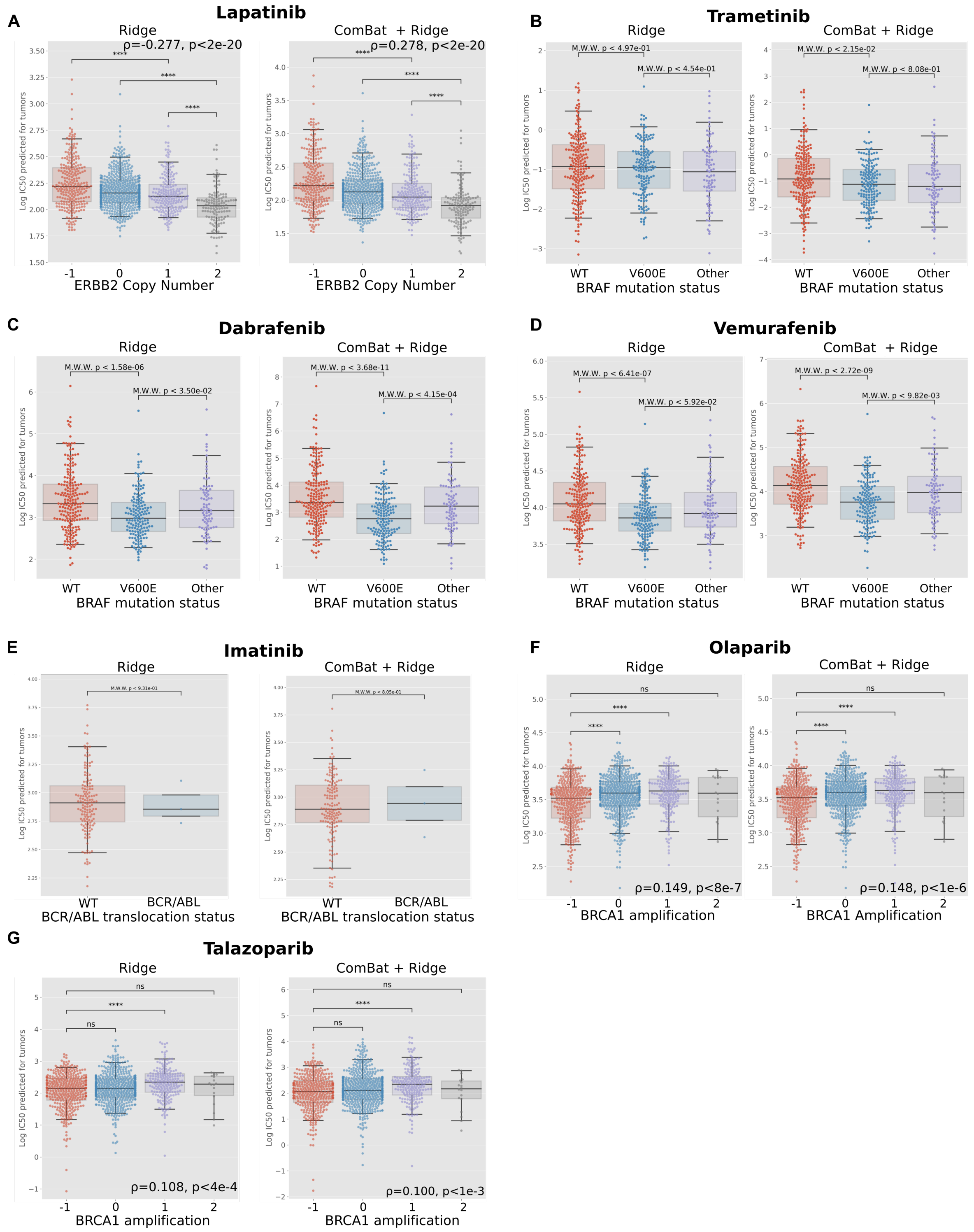


Figure Supp10: **Stratification with a Ridge regression on the bulk set of genes or with ComBat as domain adaptation.** We compared the results of Fig 4 and Fig Supp9 to two scenarios: one without any domain adaptation between cell lines and tumors, and one with ComBat as the domain adaptation step. **(A)** Lapatinib predicted response correlation with ERBB2 amplification is comparable to PRECISE, whether ComBat is used or not. **(B)** Trametinib sensitivity to $BRAF^{V600E}$ mutation, however, is not predicted. When using ComBat, a slight discrimination is observed between wild type and mutated tumors but the regression model fails to discriminates between V600E and other mutations. In Dabrafenib **(C)** and Vemurafenib **(D)**, Ridge regression and ComBat successfully indicate the sensitivity to $BRAF^{V600E}$ mutation, but the power is lower than PRECISE. BCR/ABL is not discriminated by neither Ridge nor ComBat + Ridge **(E)**. Finally, PARP inhibitors Olaparib **(F)** and Talazoparib **(G)** are also recovered, but with correlations two to three times lower than with PRECISE.

### References


Dillies, M.-A., Rau, A., Aubert, J., Hennequet-Antier, C., Jeanmougin, M., Servant, N., Keime, C., Marot, G., Castel, D., Estelle, J., *et al.* (2013). A comprehensive evaluation of normalization methods for illumina high-throughput rna sequencing data analysis. *Briefings in bioinformatics*, **14**(6), 671–683.

Geeleher, P., Cox, N. J., and Huang, R. S. (2014). Clinical drug response can be predicted using baseline gene expression levels and in vitro drug sensitivity in cell lines. *Genome Biology*.

Gong, B., Shi, Y., Sha, F., and Grauman, K. (2012). Geodesic flow kernel for unsupervised domain adaptation. In *Computer Vision and Pattern Recognition (CVPR), 2012 IEEE Conference on*, pages 2066–2073. IEEE.

Gopalan, R., Li, R., and Chellappa, R. (2011). Domain adaptation for object recognition: An unsupervised approach. In *Computer Vision (ICCV), 2011 IEEE International Conference on*, pages 999–1006. IEEE.

Robinson, M. D. and Oshlack, A. (2010). A scaling normalization method for differential expression analysis of rna-seq data. *Genome biology*, **11**(3), R25.

Zwiener, I., Frisch, B., and Binder, H. (2014). Transforming rna-seq data to improve the performance of prognostic gene signatures. *PloS one*, **9**(1), e85150.
